# Visual interpretability of image-based classification models by generative latent space disentanglement applied to in vitro fertilization

**DOI:** 10.1101/2023.11.15.566968

**Authors:** Oded Rotem, Tamar Schwartz, Ron Maor, Yishay Tauber, Maya Tsarfati Shapiro, Marcos Meseguer, Daniella Gilboa, Daniel S. Seidman, Assaf Zaritsky

## Abstract

The success of deep learning in identifying complex patterns exceeding human intuition comes at the cost of interpretability. Non-linear entanglement of image features makes deep learning a “black box” lacking human meaningful explanations for the models’ decision. We present DISCOVER, a generative model designed to discover the underlying visual properties driving image-based classification models. DISCOVER learns disentangled latent representations, where each latent feature encodes a unique classification-driving visual property. This design enables “human-in-the-loop” interpretation by generating disentangled exaggerated counterfactual explanations. We apply DISCOVER to interpret classification of in-vitro fertilization embryo morphology quality. We quantitatively and systematically confirm the interpretation of known embryo properties, discover properties without previous explicit measurements, and quantitatively determine and empirically verify the classification decision of specific embryo instances. We show that DISCOVER provides human-interpretable understanding of “black-box” classification models, proposes hypotheses to decipher underlying biomedical mechanisms, and provides transparency for the classification of individual predictions.

## Introduction

With the rapid growing volume and complexity of modern biomedical visual data, we can no longer rely on human capacity to identify visual patterns in biomedical images. Deep learning models, specifically convolutional neural networks (CNNs), have shown great promise in identifying complex patterns in biomedical images. CNNs may achieve performance comparable and even superior to that of domain experts, as shown for example in diabetic retinopathy (Gulshan et al. 2016, Ting et al. 2017, Hacisoftaoglu et al. 2020, Ruamviboonsuk et al. 2022), skin cancer (Esteva et al. 2017, <u>Fujisawa et al. 2018</u>), cardiovascular risk factors (Poplin et al. 2018), chest radiograph interpretation (Rajpurkar et al. 2018), breast cancer (Rodriguez-Ruiz et al. 2019), mesothelioma (Courtiol et al. 2019), genetic disorders (Gurovich et al. 2019), and COVID (Wang et al. 2021). While classical machine learning relies on hand-crafted features, the success of deep learning stems from data-driven nonlinear optimization of feature extraction toward a specific classification task, without relying on prior assumptions about the image data or specific measurables. However, this success comes at the cost of poor interpretability. In classical machine learning, hand-crafted features can be back-tracked to provide interpretable explanations of the model decisions (e.g., SHAP, Lundberg et al. 2017). However, CNNs’ nonlinear entanglement of image features makes deep learning a “black box” that lacks straightforward explanations. Understanding the image properties underlying the models’ prediction is especially critical in biomedical domains because the clinician/researcher must understand the clinical/phenotypic basis of the machine’s prediction in order to trust it (Belthangady et al. 2019, Andrews et al. 2022, Rajpurkar et al. 2022). Moreover, understanding the reason behind a machine’s prediction is key for deciphering the underlying biological mechanisms, which in cases of disease detection, is a critical step toward treatment.

The most common visual interpretability methods for deep learning image-based classification models are attribution-based (also known as gradient-based) methods that generate heatmaps or “attention maps” that highlight the image regions contributing most to the models’ prediction (Zhou et al. 2016, Selvaraju et al. 2017, Shrikumar et al. 2017). Another, more recent approach for visual interpretability, known as “counterfactual explanations” (e.g., Lang et al. 2021), is based on the use of generative models that alter the image to affect the model’s prediction. This is done, for example, by generating counterfactual images where the classification-driven image properties are exaggerated to enable identification of subtle phenotypes (Zaritsky et al. 2021). Alterations in image patterns that are associated with changes in the model’s prediction can then be interpreted by experts to establish new mechanistic hypotheses and draw biological or clinical conclusions (e.g., Zaritsky et al. 2021). Practically, however, current interpretability methods suffer from limitations that make them not sufficiently robust for systematic general-purpose visual interpretability of biomedical imaging based deep learning classification models (Rodríguez et al. 2021, Rudin et al. 2019). A major limitation toward systematic interpretability is the entanglement of multiple classification-driving image properties producing convoluted visual explanations of the object that is being interpreted. This hampers the expert’s ability to interpret which semantic image properties contributed to the classifier’s decision.

Here, we present *DISentangled COunterfactual Visual interpretER* (*DISCOVER)*, a generalized method toward systematic visual interpretability of image-based classification models. The main innovation of DISCOVER is a disentangling module that forces each latent feature to encode exclusive image property that is distinct from the ones encoded by other latent features, and thus, leads to disentanglement of the latent representation in the context of the image space. This disentanglement allows visually intuitive traversal of the latent space one latent feature at a time under the assumption that each feature will encode independent classification-driving semantic image properties. We demonstrated that latent features can be visually interpreted, by domain experts, to specific semantic image properties. These interpreted latent features can discover and quantify classification-driving semantic properties that did not have explicit measurements, and to rank the importance of each semantic property on instance-specific model’s predictions.

We applied our visual disentangled interpreter to the domain of in vitro fertilization (IVF). In IVF, egg(s) are removed from the patient’s ovaries, fertilized, and incubated in a laboratory. One or a few embryos from the cohort are then transferred to the patient’s uterus. IVF is an ideal example of a biomedical domain where visual assessment is the key to its success. This is specifically relevant to the visual assessment of embryo quality that occurs prior to embryo selection for transfer or cryopreservation (Gardner et al. 2000, Alpha Scientists 2011). After approximately forty years of low-throughput techniques, automated live embryo imaging technique transformed IVF into a data-intensive field and led to the development of unbiased and automated methods that rely on machine learning for visual assessment of embryo quality (Raef et al. 2019, Simopoulou et al. 2018, Bormann et al. 2020, Khosravi et al. 2019, Chavez-Badiola et al. 2020, Tran et al. 2019, Chen et al. 2019, Uyar et al. 2015, Silver et al. 2020). These advances are now revolutionizing the field, with recent studies demonstrating that deep learning models can exceed clinician performance in embryo assessment (Bormann et al. 2020, Fitz et al. 2021). The high volume of standardized image-based data that are acquired in clinics around the globe, along with the complexity of the phenotypic information in embryo images, make IVF an attractive application to showcase visual interpretability. We demonstrate the ability of DISCOVER to decipher manually annotated embryo quality properties, to discover embryo quality properties that were not explicitly annotated, and to determine which quality properties were most dominant in the classification decision for specific embryos.

## Results

### Deep learning classification of blastocyst morphologic quality

The IVF process involves retrieving a cohort of oocytes, fertilizing them with sperm, and incubating them for several days in vitro. The fertilized eggs (embryos) are typically incubated until the blastulation stage of embryonic development is reached after 5 or 6 days of development (henceforth called a blastocyst). The highest quality blastocyst(s) is then transferred into the uterus for implantation. We trained a deep neural network to predict a blastocyst binary morphologic quality (i.e., high versus low quality) using a balanced training dataset consisting of 2,170 expert-annotated blastocysts images captured after 103 hours post insemination and obtained retrospectively from three clinics (Methods). An expert embryologist annotated each blastocyst image according to two of the Gardner and Schoolcraft blastocyst quality grading criteria (herein called Gardner) (Gardner et al. 1999) (Fig. 1A): (1) morphology of the inner cell mass (ICM), a compacted grouping of cells within the blastocyst that eventually form the fetus; (2) morphology of the trophectoderm (TE), a single cell layer surrounding the blastocyst periphery that eventually forms the placenta. To define binary labels, the blastocysts were defined as either ‘high’ (N = 1,085) or ‘low’ (N =1,085) quality, based on their ICM and TE annotations, according to the criteria defined in (Gardner et al. 1999, Khosravi et al. 2019) (Methods) (Fig. 1B). We developed a preprocessing pipeline to localize blastocysts within the image (Fig. S1), followed by fine-tuning a pre-trained VGG-19 (Simonyan et al. 2014) deep convolutional neural network model by re-training it to discriminate between high-versus low-quality blastocysts (Methods) (Fig. 1C). This IVF-CLF model performed well with an area under the receiver operating characteristic (ROC) curve (AUC) of 0.93 (Fig. 1D). The classification of high-versus low-quality blastocysts was previously solved by others, with comparable results (e.g., Khosravi et al. 2019). The reason for working with a high-performing model that is based on known morphologic properties is that it allows for a controlled test-bed for assessing our interpretability method. We attempted to interpret our IVF-CLF model by applying GradCAM, a classic “explainable AI’’ method that generates heatmaps highlighting the image regions contributing most to a given prediction of deep neural network classifiers (Selvaraju et al. 2017). But GradCAM provided convoluted visual explanations that were unintuitive to embryologists (Fig. 1E).

**Figure 1.**
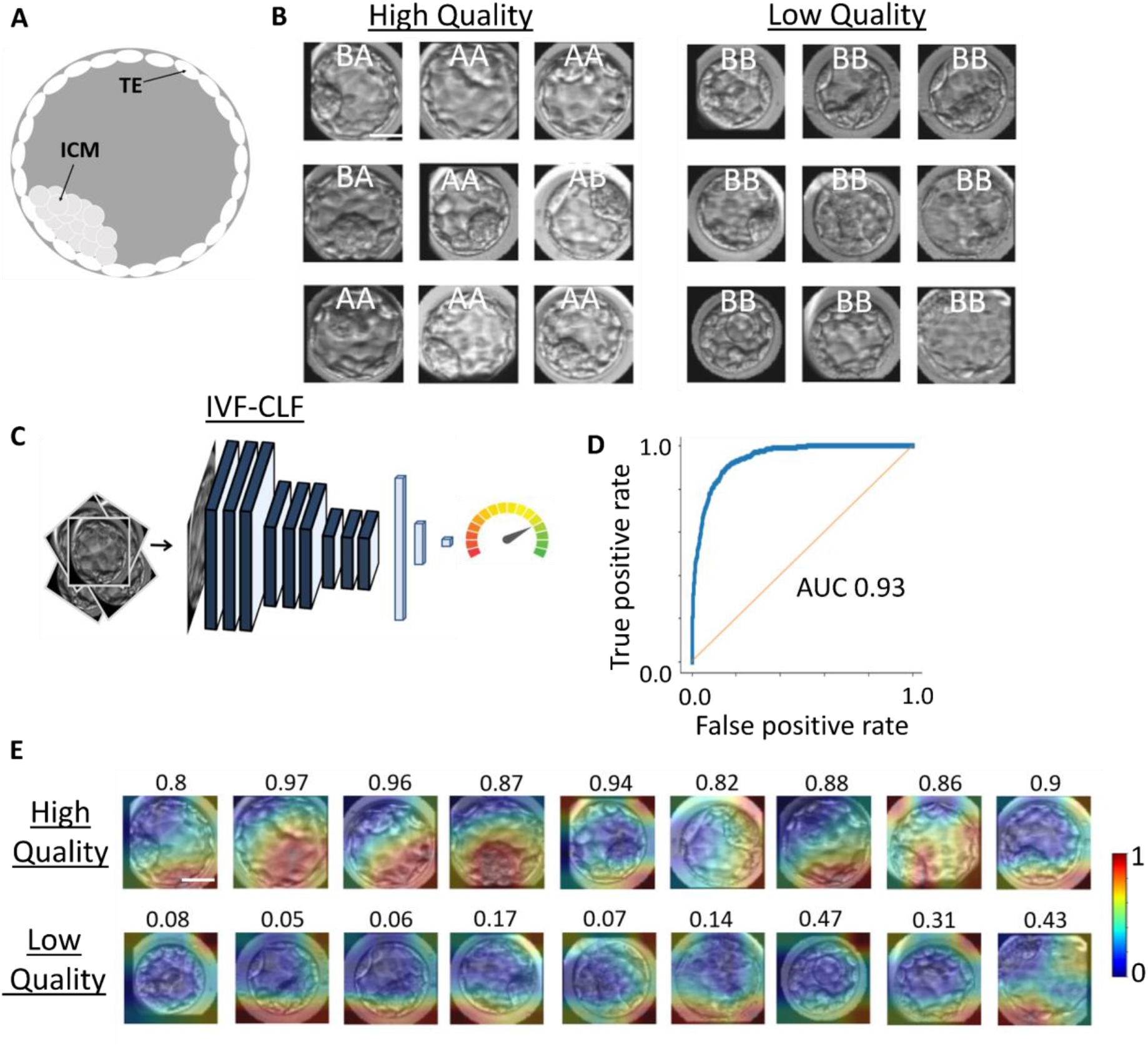
Supervised machine learning model accurately classifies blastocysts according to their high versus low morphologic quality. (**A**) Blastocyst quality is determined according to two manually annotated quality criteria, the Inner Cell Mass (ICM) and the Trophectoderm (TE). The morphologic quality of the ICM is graded A-C and determined according to the size and compaction of the mass of cells that eventually form the fetus. The morphologic quality of the TE is graded A-C and determined according to the number of cells and cohesiveness of the single layer of cells at the outer edge of the blastocyst that eventually forms the placenta. (**B**) Left: representative blastocysts labeled as high quality according to manual embryologists’ (ICM, TE) annotations of (A, A), (A, B), or (B, A) (top row). Right: Representative blastocysts labeled as low quality according to manual embryologists’ (ICM, TE) annotations of (B, B), (C, B), or (B, C). (**C**) Schematic sketch of the IVF-CLF binary classifier trained to predict the quality score of a blastocyst image [0-1]. The IVF-CLF backbone is a VGG-19 architecture and training was initialized from the ImageNet pretrained weights. 977 high-quality and 977 low quality blastocysts were used for training. (**D**) ROC curve of the blastocysts quality IVF-CLF with a test set of 108 high-quality and 108 low quality blastocysts. (**E**) GradCAM heatmaps obtained by aggregation of the last convolutional layer of IVF-CLF for all blastocysts examples in (B). Warmer colors correspond to more relevant regions for the classification outcome. For all panels scale bar = 12.5 *μ*m.

### DISCOVER, the visual disentangled interpreter - a generative network architecture for visual interpretability of image-based deep learning classification models

We developed *DISCOVER*, a general-purpose interpretability method designed to discover the underlying visual properties driving a classification task, and applied it to identify the visual cues driving the IVF-CLF trained to discriminate between high- and low-quality blastocyst images. DISCOVER is based on a deep learning generative framework that encodes the image data to a disentangled latent representation. This allowed for traversing over the latent space, one latent feature at a time, by forcing each latent feature to encode independent classification-driving image properties. This amplification of a specific discriminative latent feature enabled interpreting images with visual counterfactual explanations along a specific phenotypic axis in the image space. Enhanced interpretability was enabled by exaggerating classification-driving latent features (and their corresponding image properties), while maintaining the rest of the features (and their corresponding image properties) fixed. Training simultaneously optimizes six loss terms described below (Fig. 2A-B, full details in Methods). The weights for each loss term were optimized during training by assigning higher weights to loss terms that did not converge.

**Figure 2.**
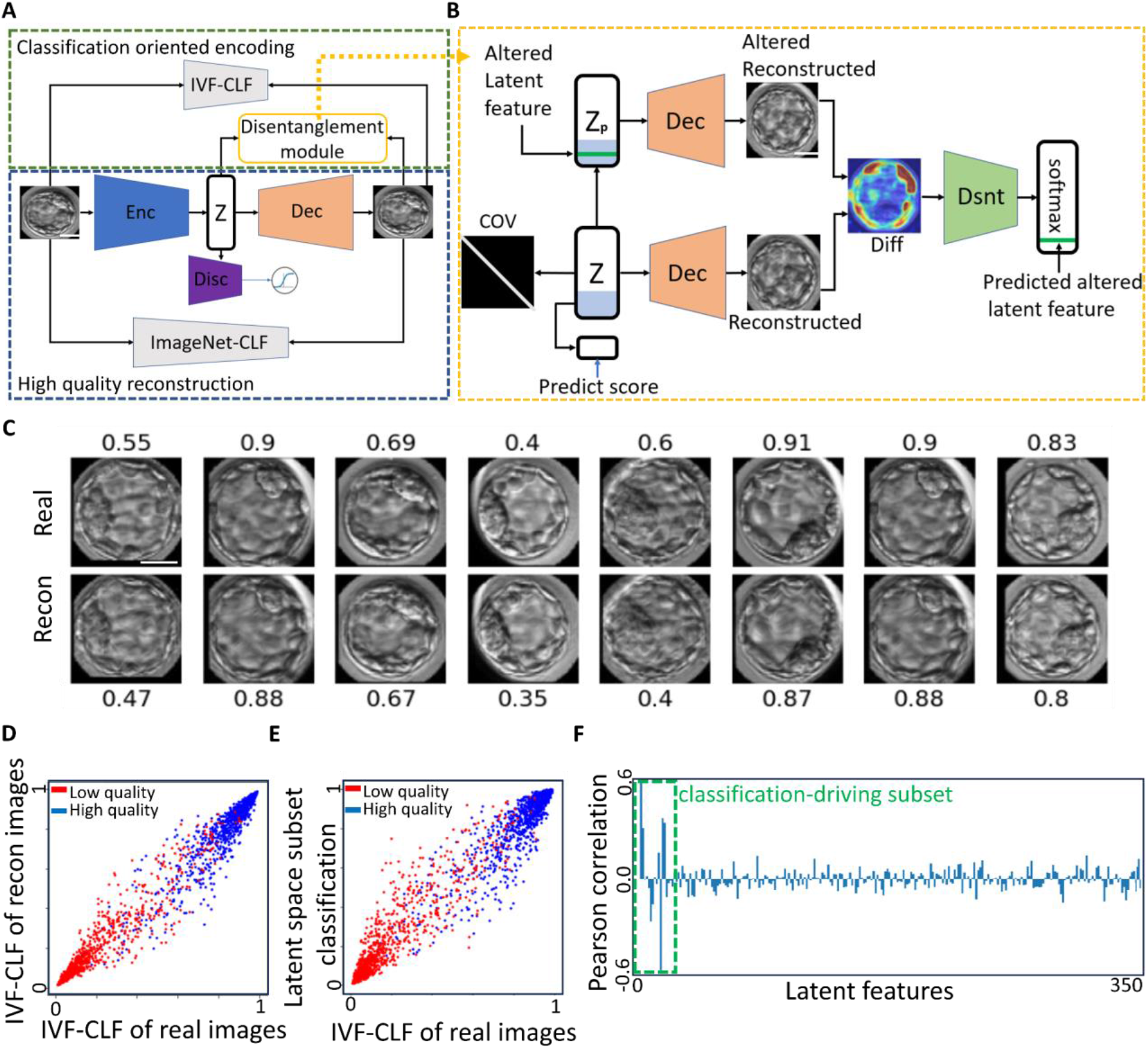
DISCOVER - a generative model designed toward visual interpretability of image-based binary classification models. (**A**) DISCOVER’s high-level architecture. Input: pre-processed blastocyst images, IVF-CLF - a binary classifier trained to predict blastocyst quality. The DISCOVER architecture is composed of 3 modules: (1) an adversarial autoencoder for high quality reconstruction and generation of realistic images from the latent representation space (dashed blue). The pre-trained ImageNet-CLF is used for perceptual loss minimization between real and reconstructed images; (2) minimization of the deviation between the IVF-CLF scores of the input image and its corresponding reconstructed image toward classification-oriented encoding (dashed green); (3) a disentanglement module (yellow, detailed in B) which decorrelates the latent features and associates a small subset of the latent features to unique image properties correlated with the IVF-CLF. Scale bar = 12.5 *μ*m. (**B**) Architecture of the disentanglement module that include two loss terms toward a classification-driving subset of latent features: (1) The disentanglement loss term minimizes the error of a new model trained to identify which latent feature was altered. This model receives as input the difference image between the unaltered reconstructed image (from Z) and the reconstructed image after altering a random latent feature (green in Zp), and is optimized to predict the index of the altered latent feature (green “predicted latent feature”); (2) Constraining the generative model to maintain a specific subset of latent features that are correlated to the frozen model’s classification score. The first 14 features in the latent representation (cyan in Z) are used as input for a new supervised model that is optimized to predict the IVF-CLF score (“predict score” below Z). Specifically, this subset of latent features is fully connected to a single neuron which is passed through a sigmoid activation, and is minimized by ‘binary cross entropy’ loss. An additional regularizer on the latent vector Z further forces decorrelation by whitening the covariance matrix. (**C**) Representative blastocysts images (‘Real’, top) and their corresponding reconstructions (‘Recon’ bottom) along with the corresponding IVF-CLF classification scores above each image. Scale bar = 12.5 *μ*m. (**D-E**) Scatter plot of the IVF-CLF classification scores of the blastocysts’ images (x-axis) and their matched reconstructed images (D) or matched scores derived from the classification-driving subset of latent features (E) (y-axis). N = 1085 high quality blastocysts (blue), N = 1084 low quality blastocysts (blue). Mean absolute error between real images scores and reconstructed images scores (D) or subset of latent features scores (E) is 0.04 and 0.06, respectively. (**F**) Pearson correlation coefficient (y-axis) between each latent feature (x-axis) and the IVF-CLF’s classification score. Panels D-F use N = 2169 blastocysts that were not used to train the model. Mean (std) of the absolute correlation of the 14 classification-driving subset of latent features were 0.257 (0.111) and 0.049 (0.0038) for the rest of the latent features. Mann–Whitney-U test p-value < 0.003.

To enable effective visual interpretability with counterfactual examples, the generative model must support reconstruction of high quality, realistic images from the latent representation space. We trained an adversarial perceptual autoencoder comprising two loss terms. The first loss term was a perceptual loss that enforced high quality image reconstruction. It was implemented by an autoencoder with a latent representation of 350-dimensions, where the reconstruction minimized the Euclidean distance between feature maps extracted from an ImageNet-based pre-trained VGG-19 network (Imagenet-CLF). This perceptual loss was previously shown to improve image embeddings (Pihlgren et al. 2020). The second loss term was an adversarial loss that enforced a continuous and probabilistic latent space. The adversarial loss optimized the latent representations such that a discriminator network fails to distinguish the latent representations derived from blastocysts images from vectors drawn from the latent space. Together, the perceptual adversarial autoencoder enabled reconstruction of realistic blastocyst images from traversals over the latent space, as validated by a trained embryologist (Fig. 2C).

The third loss enforced domain-specific classification-oriented encoding. Subtle differences in visual features important for the supervised model’s decision may be lost during image reconstruction. Thus, we minimized the discrepancy of the supervised model’s intermediate layers (i.e., perceptual loss) and the IVF-CLF prediction score between the input images and their corresponding reconstructed images. This second perceptual loss constrains the generative model to maintain image features that are important for the supervised model’s decision. Accordingly, the blastocysts images and their corresponding reconstructions exhibited similar IVF-CLF classification scores (Fig. 2C-D).

The fourth and fifth loss terms enforced disentanglement of the latent representation (Fig. 2A, yellow and Fig. 2B). The goal of these loss terms was to constrain a latent representation such that each latent feature encodes a distinct visual property in the image. This disentanglement was achieved by (1) whitening (forth loss) by decorrelating the latent space, and forcing its covariance toward a unit matrix (Bardes et al. 2021) (Fig. 2B, ‘COV’ matrix, Fig. S2), and (2) counterfactual disentanglement (fifth loss) by optimizing a new network (Fig. 2B, green ‘Dsnt’ trapeze) to identify which latent feature was altered in a perturbed image. The input of the counterfactual disentanglement model consisted of two images: the unaltered reconstructed image and the reconstructed image after altering the latent feature (Fig. 2B). These two loss terms constrain each latent feature to encode image features that are distinct from other latent features and, thus, leads to disentanglement of the latent representation. This allows for simpler traversal of the latent space one feature at a time under the assumption that each feature will encode independent classification-driving image features. We also hypothesize that such feature disentanglement will push the latent representation, such that each latent feature will tend to encode a single image feature. In summary, the disentangled latent representation enables more intuitive visual interpretability where alteration of each latent feature would amplify image properties specifically assigned to that feature. This is in contrast to entangled latent representations, where each latent feature is more prone to encode uninterpretable visual image properties.

The sixth, and final, loss term, enforced a classification-driving subset of latent features (Fig. 2B, cyan feature subset marked in Z). The goal of this loss term was to attain a sub-group of latent features that are highly correlated to the classification model’s prediction, while the rest of the latent features maintain high quality reconstruction. We forced 14, out of the 350 latent features, to correlate more strongly with the classification output of the input image. This was achieved by (simultaneously) training another layer (of a single neuron) to predict the IVF-CLF’s classification score from the first 14 features in the latent representation. Accordingly, the IVF-CLF’s classification scores were highly associated with the corresponding classification derived from the 14-dimensional subset (Fig. 2E). These first 14 latent features were more correlated to the IVF-CLF classification score when compared to the other features in the latent representation (Fig. 2F).

All six loss terms were minimized simultaneously, ultimately providing us with a generative model designed for interpretation and discovery of blastocyst quality classification-driving clinically meaningful image properties. Specifically, a generative model enabling high-quality and realistic reconstruction (loss #1) and traversal (loss #2) of the latent space, with a domain-specific classification oriented encoding (loss #3). The latent representation included a subset of 14 latent features optimized toward explainability by visual disentanglement (loss #4-5) and correlation with the classifier that is being interpreted (loss #6).

Ablation experiments verified that all loss terms were necessary toward high quality reconstruction (Fig. S4A), classification oriented encoding (Fig. S4B), classification-driving subset of latent features (Fig. S4C-D), and disentanglement of the latent representation (Fig. S4E).

### Visual interpretation of classification-driving latent features: blastocyst size and trophectoderm

To visually interpret which blastocyst morphologic quality properties had the greatest impact on the classification, we ranked the subset of classification-driving latent features according to their correlations with the IVF-CLF’s classification score. For each of the top ranked latent features and for each given blastocyst, we generated a series of counterfactual explanations. By decreasing and increasing each current latent feature by 3 standard deviations, while fixing all other features, the decoder could generate a series of ‘‘in silico’’ blastocysts images gradually morphing toward exaggerated better or worse quality along the visual phenotypic axis defined by that feature, in accordance with the IVF-CLF’s classification score (Fig. 3A, Fig. S5A-B). We visualized the counterfactual visual alteration for each of the top five ranked features of the same reconstructed blastocyst image. The visualization of the counterfactual alteration was computed using the Structural Similarity index (SSIM) (Renieblas et al. 2017), where each pixel was assigned with the SSIM dissimilarity of its corresponding patch between two reconstructed images (Methods). Visualizing each feature in respect to reconstructed images after major alterations (± 3 standard deviations), for the same blastocyst, revealed that each feature showed a distinct visual counterfactual alteration pattern (Fig. 3B). These results suggested that the classification-driving latent features were visually disentangled by the morphologic properties that they encode in the reconstructed blastocyst images.

**Figure 3.**
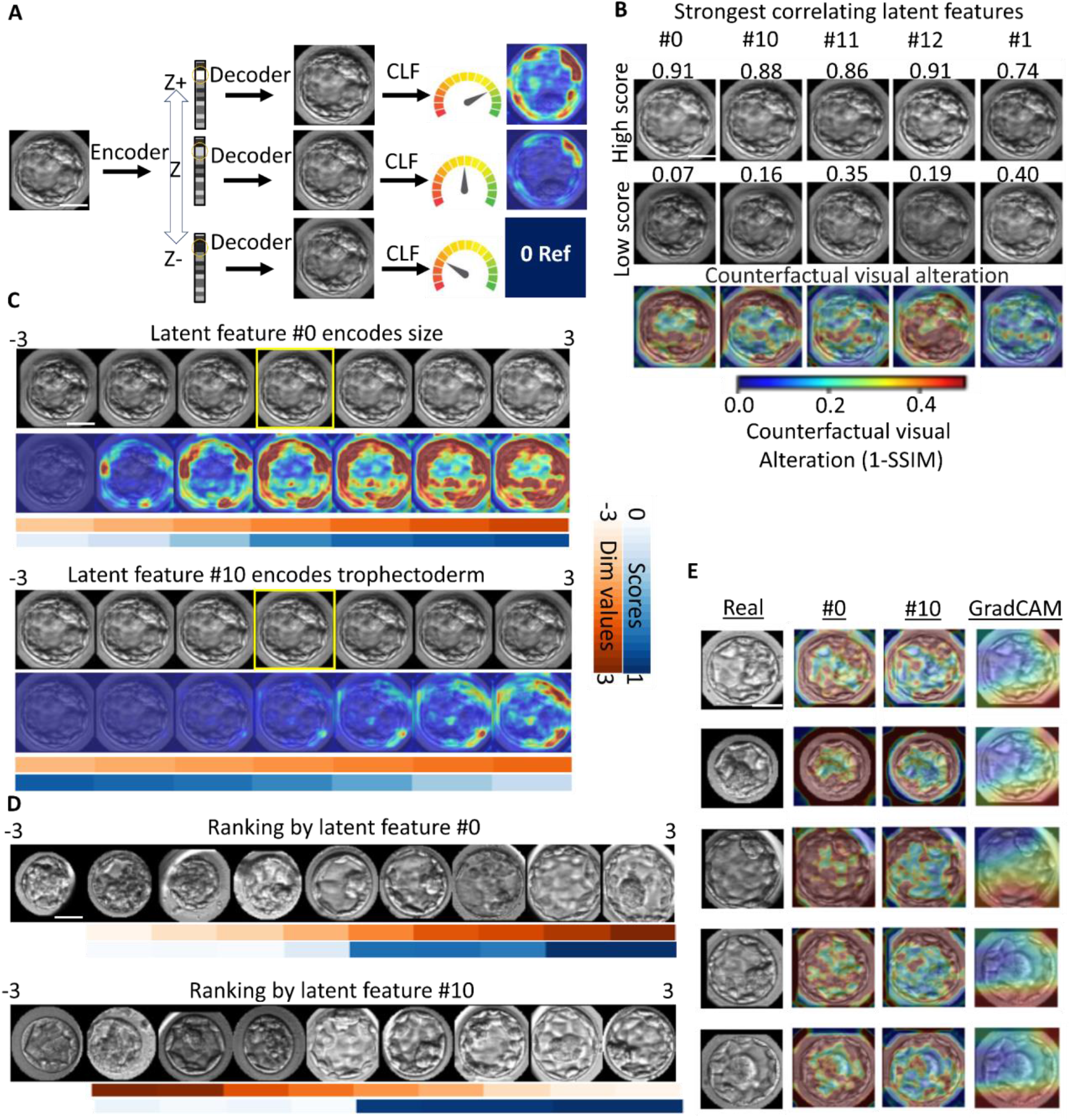
Visual interpretability by DISCOVER identifies blastocysts’ size and TE quality as classification-driving image properties encoded by the top two features in the latent representation. (**A**) Approach: Visual interpretability via counterfactual explanation with DISCOVER. From left to right. A blastocyst image is encoded to its corresponding latent representation. A latent feature is gradually altered while fixing all other features in the latent representation. The altered latent representations are decoded to their corresponding reconstructed blastocysts images. The reconstructed blastocysts sequence can be validated according to a gradual change in their corresponding classifier score and interpreted according to visualization of their counterfactual visual alteration. (**B**) Counterfactual visual alteration of the same blastocyst according to the alteration (± 3 standard deviations) of the five latent features most correlated to the IVF-CLF, left-to-right in descending order (Pearson correlation coefficient): #0 (0.69), #10 (−0.65), #11 (0.44), #12 (0.4), #1 (0.36). Top row: reconstructed altered images with increased classification score. Middle row: reconstructed altered images with reduced classification scores. Bottom row: counterfactual visual alteration between the corresponding top and middle rows. Color map indicates the local change measured as 1-SSIM, and is also used in panels C and E. (**C**) Gradual traversal of the two latent features #0 and #10 that were the most correlated to the classifier score. Traversal was performed in the range of −3 and 3 standard deviations around the original encoded value. Top: reconstructed images. Bottom: counterfactual visual alteration. Red - latent feature values, blue - classification scores. Yellow bounding box - reconstruction of the unaltered image. (**D**) Panel of nine randomly selected sequences of blastocysts in predefined monotonically increasing intervals of latent feature #0 (top) and #10 (bottom). Red - latent feature value, blue - classification score. (**E**) Comparison of DISCOVER interpretability to GRADCAM. Five examples showing (from left to right): the original blastocyst, visual counterfactual alteration of latent feature #0 and #10, and GradCAM heatmap obtained by aggregation of the last convolutional layer. For all panels scale bar = 12.5 *μ*m.

We next used these visual counterfactual alterations to interpret the two top classification features. These were features #0 and #10 with a Pearson correlation coefficient of 0.69 and −0.65 to the IVF-CLF, respectively. Since the variance of all latent features equals one due to the latent generative-adversarial loss (Methods), we morphed the latent features within the range [−3, +3], and visualized the counterfactual alterations between the two extreme reconstructed images. We observed that the counterfactual visual alterations of feature #0 were concentrated around the blastocyst bulk, indicating a monotonically altered blastocyst size, leading to a corresponding change in the classification score (Fig. 3C - top, Fig. S5B-C). While the blastocyst size was not explicitly annotated in our data, it was previously linked to clinical pregnancy (Sciorio et al. 2021). The blastocyst size is also a property highly associated with the blastocyst expansion status (Lagalla et al. 2015), i.e., the volume and degree of expansion of the blastocyst cavity, which is the third quality grading criteria in the Gardner assessment (Gardner et al. 2000). For feature #10 we observed visual counterfactual alterations concentrating in the blastocyst periphery, which corresponds to the trophectoderm. The counterfactual trophectoderm visual quality was monotonically altered in concurrence with the latent feature value, leading to a corresponding change in the classification score (Fig. 3C - bottom, Fig. S5B-C). These visual explanations for latent features #0 and #10 were robust to image flipping and brightness changes (Fig. S5D). To further corroborate the encoding to blastocyst size and TE quality, we randomly selected a sequence of nine blastocysts in predefined monotonically increasing intervals of latent features #0 and #10. Visual observation by embryologists suggested that the changes were mostly attributed to blastocyst size and TE quality, and respectively, the IVF-CLF scores gradually increased in relation to the change in the corresponding latent features (Fig. 3D, Methods). These disentangled visual explanations of size and TE could not be attained with GradCAM (Fig. 3E). These results established the potential for DISCOVER to generate representations in which each latent feature encodes a visually interpretable classification-driving image property.

### Quantitative and empirical expert validation of interpreted classification-driving latent features encoding the blastocyst size and the trophectoderm

After visually interpreting the classification-driving latent features #0 and #10 as blastocyst size and trophectoderm, correspondingly, we aimed at quantitatively and systematically validating these interpretations. Correlation between the latent features showed that features #0 and #10 were weakly correlated (Pearson correlation coefficient = −0.35, ranked 1 out of 91 pairwise feature correlation, see red dashed square in Fig. S2). Moreover, it is known that the blastocyst size and TE quality are associated with one another and with the overall blastocyst quality (Lagalla et al. 2015). To overcome the challenge of quantitatively decoupling the interpretation of these associated latent features to their corresponding associated morphologic properties, we matched pairs of blastocysts such that one morphological property (size/TE) was similar among the blastocysts and the other property was different. Specifically, to quantify the association between latent feature #0 and blastocyst size, we matched pairs of blastocysts with the same expert embryologist-annotated TE grades (both grade ‘A’ or both ‘B’), and with large differences in their sizes, as calculated from the segmentation masks. Such matching enabled direct comparison of size by reducing the confounding effect introduced by the correlated TE. To assess the association between latent feature #0 and blastocyst size, we calculated the distribution of signed differences in feature #0 between the larger and the smaller blastocysts in the matched pairs. Most of the larger blastocysts in the matched pairs had higher values in feature #0 as observed by a distribution shifted toward higher positive values (Fig. 4B, blue distribution), indicating that larger blastocysts (with the same TE annotations) were associated with higher values in latent feature #0. As a control, we calculated the distribution of signed differences of latent feature #10 in the matched pairs. Here, we flipped the order of subtraction because feature #10 was negatively correlated with the IVF-CLF scores. This distribution was mostly centered around 0 indicating that latent feature #10 was only marginally altered for larger blastocysts with matched TE annotations (Fig. 4B, red distribution). This direct comparison between distributions was legitimate because the latent features were normalized and indicated that latent feature #0 was more associated with the blastocyst size. To further validate that blastocyst size was specifically controlled by feature #0, we repeated the process of calculating the distributions of the matched blastocysts pairs’ signed differences for each of the 14 classification-driving subsets of latent features (Methods). The subtraction order was according to the correlation sign of each latent feature with the IVF-CLF scores (Fig. 2F). The median of the differences between larger versus smaller blastocysts pairs with matched TE annotations was highest for feature #0, thereby providing more evidence that this feature specifically encodes the blastocyst size (Fig. 4C).

**Figure 4.**
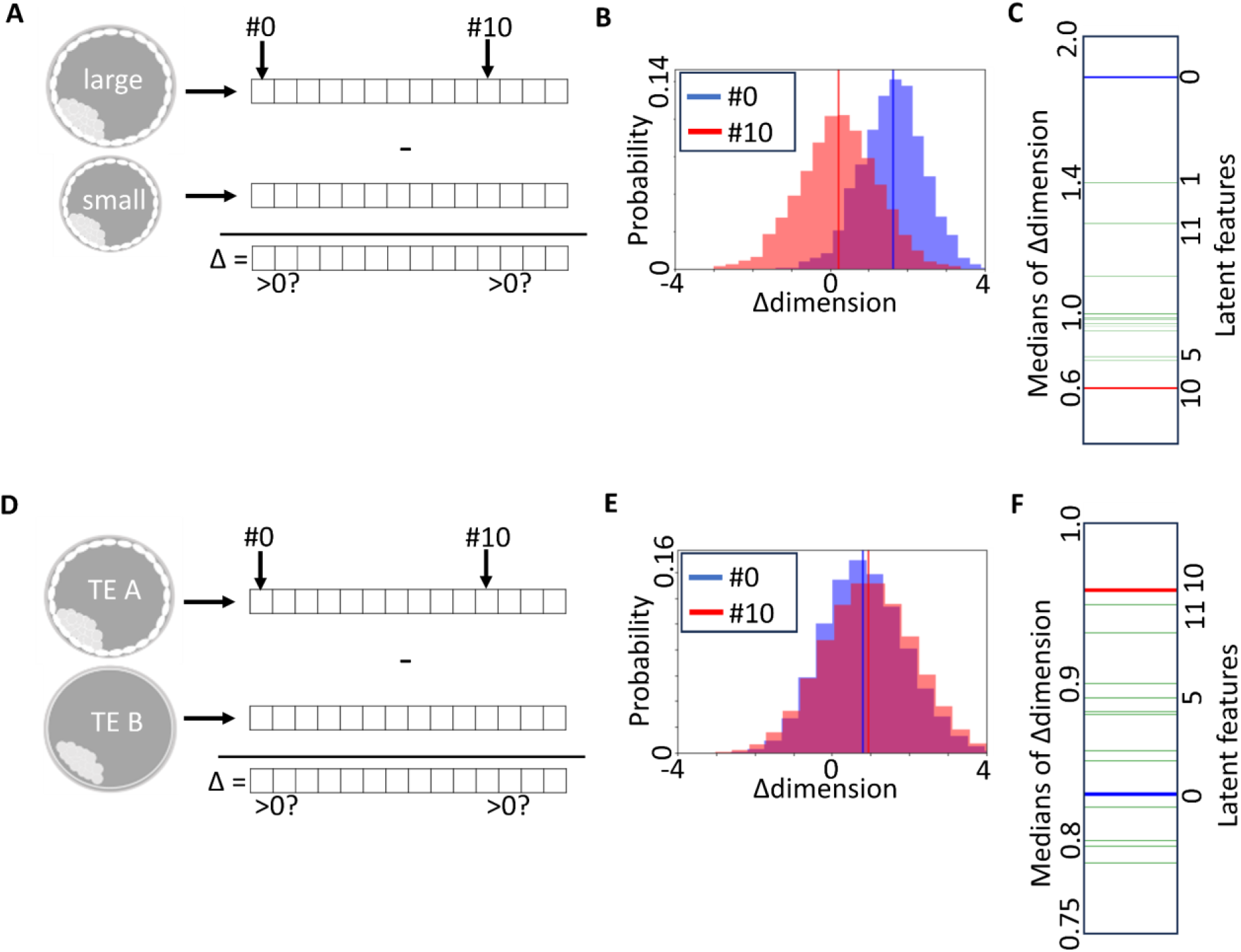
Statistical validation that the blastocysts’ size and TE quality are encoded by the top two features in the latent representation. (**A**) 2,134 matched pairs of blastocysts with similar TE annotations and different sizes. The subtractions of each latent feature value for each blastocyst and its corresponding paired blastocyst were pooled for each latent feature. The order of subtraction is determined according to the blastocysts’ size and sign of the correlation between the latent feature and the IVF-CLF scores (Fig. 2F). (**B**) Distributions of signed differences in latent features #0 (blue) and #10 (red) between matched pairs of blastocysts with similar TE and different size. Median values = 1.64 and 0.23 respectively (vertical lines). (**C**) Median values of the distributions of signed differences for the 14 classification-driving subsets of latent features. The blue and red vertical line represent the median of latent features #0 and #10 respectively. (**D-F**) Analysis of 808,326 matched pairs of blastocysts with similar size and different TE, corresponding to panels A-C. (**E**) Median values = 0.8 (latent feature #0) and 0.96 (latent feature #10). (**F**) Note smaller dynamic range in respect to C.

We repeated the same analysis to quantitatively link latent feature #10 to the TE quality. We matched pairs of blastocysts with similarly computed sizes and differently annotated TE grades (‘A’ with ‘B’ or vice versa) and calculated the distribution of signed differences in feature #10 between the blastocysts with lower and higher TE grades (Fig. 4E). Blastocysts with higher TE qualities (and similar sizes) were associated with positive difference values in latent feature #10 (Fig. 4E, red). Using latent feature #0 as a control, showed positive difference values to a lesser extent (Fig. 4E, red versus blue). The milder effect in feature #10 in respect to #0 could be caused because of imperfect segmentation of the blastocyst and/or because the imperfect disentanglement of feature #0, in terms of its phenotypic uncoupling - i.e., latent feature #0 may contain some information specifically attributed to the TE in addition to size (see Discussion). Still, latent feature #10 encoded the TE quality better than any other of the classification-driving subset of latent features (Fig. 4F).

As a final validation, we decided to empirically assess whether a trained embryologist can specifically associate the deviation in a latent feature with its corresponding interpreted morphologic property. We matched pairs of blastocysts according to latent features #0 and #10. This time, we did not use the annotated TE and computed size; rather, we aimed for expert inference of these morphologic properties from the latent features’ values. Matched blastocyst pairs had either similar values for latent feature #0 and dissimilar values for latent feature #10 or vice versa. A trained embryologist was provided with images of each matched pair and asked to determine whether blastocysts were different in size or in TE quality, while knowing that one of these parameters was fixed (i.e., highly similar). The embryologist was able to identify the different latent features according to the corresponding interpreted morphological property in 65/75 (86%) of pairs (Fig. S3). When asked to determine for which blastocyst the TE was better in pairs that had similar values of feature #0 and dissimilar values of feature #10, the embryologist successfully identified 33/39 (85%) of blastocysts with “better” feature #10. When asked to determine for which blastocyst the size was larger in pairs that had similar values of feature #10 and dissimilar values of feature #0, the embryologist successfully identified 31/36 (86%) of blastocysts with “better” feature #0. Altogether, our results established that DISCOVER visually disentangled the latent representation, such that latent feature #0 specifically visually encodes the blastocyst’s size and latent feature #10 specifically visually encodes the trophectoderm’s quality.

### Discovery and interpretation of the blastocoel density as a classification-driving property

Our previous results established that DISCOVER can identify latent features that encode two hallmark embryo morphologic properties, according to the Gardner blastocyst assessment system: blastocyst size and TE quality. Both of these properties are routinely assessed by embryologists to determine blastocyst quality prior to implantation. Next, we asked whether we could use DISCOVER to identify latent features that encode non-obvious morphologic properties in the blastocyst, i.e., ones that were not used during manual blastocyst quality annotation? To answer this question, we turned our attention to latent feature #11, the third top classification feature (Fig. 3B) with a Pearson correlation coefficient of 0.44 in relation to the IVF-CLF score (Fig. 2F). Latent feature #11 also appeared in Fig. 4C and Fig. 4F as one of the top 3 features most correlated with blastocyst size and TE quality, which further indicates that it encodes discriminative information about the blastocyst’s quality. The visual counterfactual alteration of latent feature #11 in Fig. 3B was identified by three embryologists / IVF experts as a potentially known morphologic feature of the embryo termed the blastocoel, a fluid-filled cavity inside the blastocyst (Shahbazi et al. 2020) (Fig. 5A). The presence and degree of blastocoel expansion, i.e., the increase in blastocoel volume is associated with implantation success and live birth (Du et al. 2016). Visual counterfactual alterations were interpreted by expert embryologists as having denser and more granular blastocoelic regions, suggesting that this change in the blastocoel appearance is the classification-driving morphologic property encoded by latent feature #11 (Fig. 5B). This visualization suggests that there are additional morphologic parameters of the blastocoel beyond its volume expansion that may be associated with overall embryo quality. A sequence of nine blastocysts that were randomly selected in predefined monotonically increasing intervals of latent features #11 further verified the encoding to the blastocoel (Fig. 5C). This interpretation of a blastocyst morphologic property that was not explicitly used to annotate blastocyst quality highlights the potential for DISCOVER to define a quantitative measure for morphologic properties that do not have explicit measurements and even identify novel visual classification-driving properties that were not known a priori.

**Figure 5.**
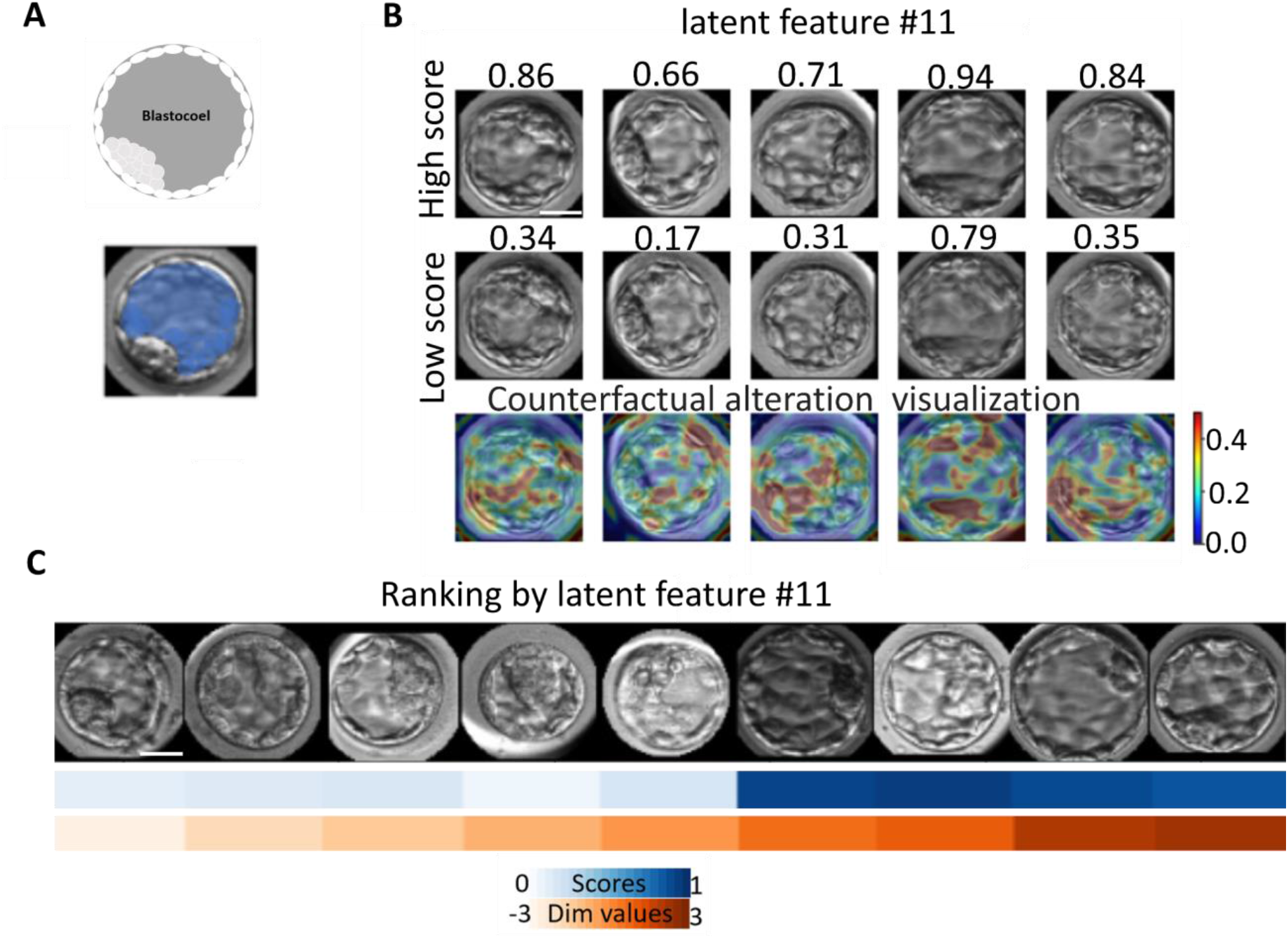
Blastocoel discovered property. (**A**) The blastocoel is a fluid-filled cavity forming the blastula marked in gray (illustration, top) and blue (blastocyst image, bottom). (**B**) Counterfactual visual alteration of five blastocysts obtained by altering latent feature #11 by ± 3 standard deviations. Top row: reconstructed altered images with increased classification scores. Middle row: reconstructed altered images with reduced classification scores. Bottom row: counterfactual visual alteration between the corresponding top and middle rows. Color map indicates the local change measured as 1-SSIM. (**C**) Panel of nine randomly selected sequences of blastocysts in predefined monotonically increasing intervals of latent feature #11. Red - latent feature value, blue - classification score. For all panels scale bar = 12.5 *μ*m.

### Determining the cause of classification of a specific blastocyst

Our results indicate that DISCOVER can reverse engineer the inner working of binary classification models by identifying classification-driving morphological properties. However, these results do not answer the question: what morphological properties drove the classification of a <u>specific</u> blastocyst? To answer this question, we took advantage of DISCOVER’s disentangled latent representation, i.e., learning representations where each latent feature is mapped to a distinct visual property in the image. This enabled us to refer to the latent representation as an (interpreted) tabular feature vector, on which we could apply SHapley Additive exPlanations (SHAP), a method for interpreting tabular-based models’ predictions (Lundberg et al. 2017). For a given prediction, SHAP calculates the contribution of each feature toward the prediction. We applied SHAP to the classification-driving subset of latent features, in the context of the prediction by the single layer perceptron model (see “predict score” in Fig. 2B) that was optimized to predict the IVF-CLF score in loss #6. The weight (“Shapely value”) attributed to each latent feature, along with the mapping from individual latent features to interpreted semantic properties, enables to identify and rank the semantic properties most influencing the classification of a specific instance (Fig. 6A). Calculating the mean SHAP values for all features across the entire dataset showed similar ranking to the correlation-based analysis with latent features #0, #10 being the two highest ranked features, and agreement in 4 of the top 5 latent features (Fig. S6). To evaluate why a specific blastocyst was predicted as high/low quality by the IVF-CLF, we visualized blastocysts according to their IVF-CLF predictions and their SHAP explanations. These visualizations were observed and described by an expert embryologist. Blastocysts with strong positive/strong negative SHAP values for feature #0 exhibited corresponding large/small sizes (Fig. 6B left), while blastocysts with strong positive/strong negative SHAP values for feature #10 exhibited corresponding high/low TE grades (Fig. 6B middle). Blastocysts with dominant positive/negative SHAP values for features #0 and #10 exhibited appropriately corresponding size and TE morphologies (Fig. 6B right). Blastocysts with strong positive SHAP values for feature #11 were confirmed to have high quality blastocoels, and were described by an expert embryologist as having high density cell regions and associated stretched zona-pellucida membranes (Fig. 6C left and middle). Blastocysts with strong negative SHAP values for feature #11 were confirmed to have low quality blastocoels (Fig. 6C right). These results indicated that SHAP can be used to weigh and rank the latent features of a specific blastocyst according to their predictive contribution, and that this ranking can be translated to the specific disentangled and interpreted morphological properties that drive the prediction of a specific blastocyst.

**Figure 6.**
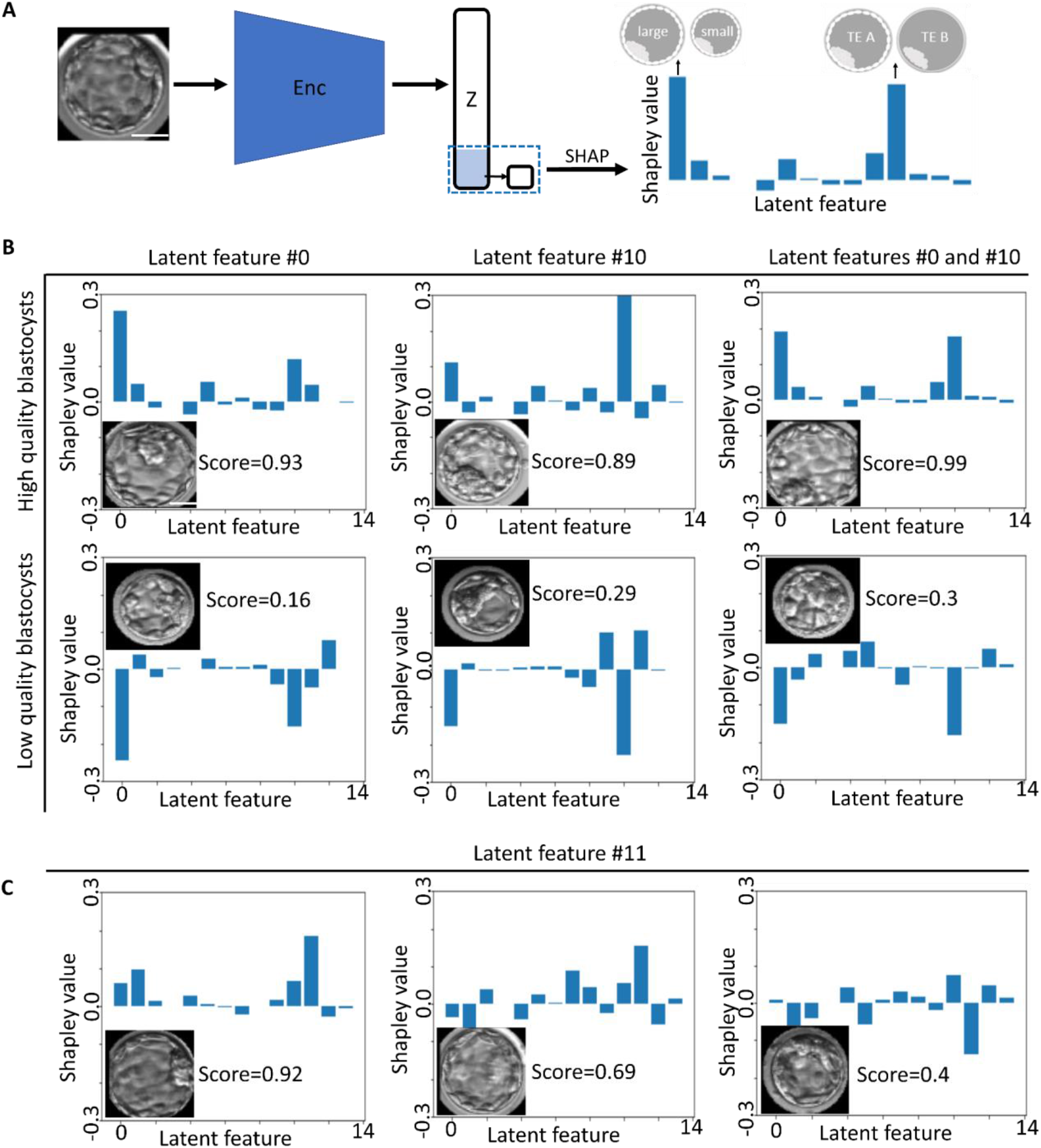
Explaining the IVF-CLF decision for a specific blastocyst by applying SHAP to the classification-driving subset of latent features. (**A**) DISCOVER’s classification-driving subset of latent features (loss #6) uses the latent representations (‘Z’) as an input to a single neuron which was trained to predict the IVF-CLF classification score. SHapley Additive exPlanations (SHAP) were applied to interpret which were the most important latent features (according to their “Shapley values”) for the prediction of this single layer perceptron given a specific instance. The interpretation of latent features to semantic properties enables instance interpretability. **(B-C)** SHAP values for specific blastocysts. (**B**) Blastocysts with dominant SHAP values for latent feature #0 (encoding size), and/or #10 (encoding TE). Top/bottom rows present IVF-CLF predicted high/low quality blastocysts correspondingly, exhibiting different explanations according to their SHAP values. Blastocysts with high SHAP values for latent feature #0 (left), high SHAP values for latent feature #10 (middle), and high SHAP values for both latent features #0 and #10 (right). **(C)** Blastocysts with dominant SHAP values for latent feature #11 (encoding the blastocoel). Shown are three blastocysts, two with high (left, middle) and one with low (right) SHAP values for latent feature #11 (our dataset had six blastocysts with the most dominant SHAP values in latent feature #11). Scale bar = 12.5 *μ*m for all panels.

### Generalizing DISCOVER to interpretation of natural images: visual interpretation of classification-driving features distinguishing between male and female facial images

We designed DISCOVER as a generalized method for visual interpretability of image-based classification models. To showcase this generalization we turned to the domain of natural images and asked whether DISCOVER can interpret the visual traits semantically distinguishing between human male and female facial images. We trained a face classifier GENDER-CLF by fine-tuning a pre-trained VGG-19 network to discriminate between male and female facial images using the celebA dataset (Liu et al. 2014) (Fig. S7A, Methods). We trained DISCOVER using the trained face classifier GENDER-CLF (identical to IVF-CLF, Methods) and we interpreted the top three ranked latent features, namely #2, #4, and #3, with Pearson correlation coefficient of 0.68, 0.49, and 0.42, respectively (Fig. S7B). Visualization of the counterfactual alteration revealed that feature #2 encoded the cheeks and jawline (smaller face for females), feature #4 the eyebrows and hair (thinner hair for females), and feature #3 the eyes (darker for females) (Fig. S7C, Methods). These traits were consistent with previous studies that highlighted cheeks, eyes and eyebrows as discriminative facial characteristics (Bannister et al. 2022, https://arxiv.org/abs/1805.00371). These results indicate that DISCOVER is a generalized interpretability method.

## Discussion

### DISCOVER is a generic framework designed toward visual interpretability of image-based classification models

Convolutional deep neural networks success at complex pattern recognition in images is attributed to non-linear simultaneous optimization of feature extraction and model training. However, this success comes with cost. The non-linear entanglement of image features makes it difficult to interpret which semantic image properties were most important for the models’ decision. DISCOVER is a generative model that optimizes latent representations geared toward interpretability of the inner decision making of a given classification model. DISCOVER representations are optimized toward classification-driven disentanglement of the latent representation, where a subset of latent features encapsulates the discriminative information of the classification model, and where each of these latent features encodes a distinct visual property in the image. Moreover, DISCOVER enables realistic reconstruction and traversal of the latent space, without losing visual information important to the classification model. Together, these design choices of DISCOVER enable expert-in-the-loop interpretation of the classification model by generating counterfactual images where each disentangled classification-driving image property is specifically exaggerated. This is achieved by shifting the latent representations and their corresponding image reconstructions, one latent feature at a time, while leaving the rest of the latent representation fixed. This counterfactual traversal along the latent space provides critical insight regarding which semantic image properties are most important for the classification model’s decision process, including discovery of new potential classification-driving semantic properties that were not known a priori. Once latent features are visually interpreted to specific semantic image properties, standard tabular-based explainable AI methods (e.g., SHAP), can be applied to weight and rank the semantic properties most influencing the classification of a specific instance. Altogether, our general framework proposes a new two-step interpretability approach. First, domain experts interpret the specific classification-driving semantic image properties encapsulated in DISCOVER’s latent representation, revealing the inner workings of a classification model. Second, using this mapping, from a latent feature to a semantic property, to explain the classification decision of specific instances. Demonstrating applicability to one biomedical (IVF) and another general computer vision (faces) datasets suggest that DISCOVER is a generalized interpretability method.

### DISCOVER interpretation of in vitro fertilization blastocysts quality classification

Our main demonstration of the applicability of DISCOVER was in the challenging domain of biomedical imaging, where providing insight explaining the “black box” prediction can propose new hypotheses to decipher the underlying biomedical mechanisms and/or assist in clinical decisions. Specifically, we interpreted a classification model optimized to predict human blastocysts morphologic quality in the context of IVF. First, we visually interpreted the top two classification-driving latent features that encode two well established blastocyst quality grading parameters: blastocyst size, as a proxy of development stage and degree of expansion, and trophectoderm quality. Second, we quantitatively and systematically validated the specific interpretation of these latent features as encoding the size and the trophectoderm, overcoming the inherent association between these two morphological properties. Third, we discovered a latent feature encoding the blastocoel density, which was a classification-driving morphological property that was not explicitly annotated. Importantly, there were no previous measurements to quantify blastocoel density, highlighting the potential of DISCOVER to discover new classification-driving semantic image properties and quantify these properties even without previous explicit measurements, through the corresponding latent feature values. Finally, we computationally determined and empirically verified which interpreted morphological properties were most important toward a classification decision of specific blastocyst instances. Our analyses demonstrate that DISCOVER can provide human-interpretable understanding of a “black-box” classification model and for the classification of individual predictions.

DISCOVER can have direct clinical relevance in the domain of IVF by providing transparency and trust in the upcoming era of “black-box” AI-based blastocyst selection (Nagaya et al. 2022, Diakiw et al. 2022, Wang et al. 2021, Sawada et al. 2021). Moreover, in situations where more than a single blastocyst is selected for transfer, the embryologist might prefer to select blastocysts with differing morphologic properties that contribute to its high-quality, under the assumption that different “mechanisms” may complement and thus increase implantation potential (and perhaps also decrease the risk of multiple pregnancy). DISCOVER was designed as a general-purpose visual interpretability of image-based classification models, and thus, can enable computational-driven biological and clinical discovery in other domains beyond IVF (Gulshan et al. 2016, Ting et al. 2017, Hacisoftaoglu et al. 2020, Ruamviboonsuk et al. 2022, Esteva et al. 2017, <u>Fujisawa et al. 2018</u>, Poplin et al. 2018, Rajpurkar et al. 2018, Rodriguez-Ruiz et al. 2019, Courtiol et al. 2019, Gurovich et al. 2019, Wang et al. 2021). Capitalizing on the AI’s unprecedented ability to automatically identify hidden semantic image patterns that are buried in complex biomedical images, along with DISCOVER’s counterfactual-based visual-guidance, there is significant potential to open the door to the generation of new biological mechanistic insight and testable hypotheses by reverse engineering machine predictions.

### DISCOVER was designed to overcome limitations of alternative image-based interpretability methods, especially toward interpretability of biomedical images

Visual interpretability methods for deep learning image-based classification models can be categorized under two broad strategies, attribution based and counterfactual based. Attribution based methods compute saliency maps, indicating how much each pixel contributed to the prediction (Ribeiro et al. 2016, <u>Zhou et al. 2019</u>, Selvaraju et al. 2017, Chattopadhyay et al. 2017, Ramaswamy et al. 2020, Ali et al. 2021). This is achieved by computing the attention of inner layers of the model by aggregating their activations, or gradients, for each pixel (Bach et al. 2015, Achtibat et al. 2023, Gur et al. 2021). Accordingly, saliency maps visualize localized regions particularly important for the classification. Such approaches are not suitable when the classification-driving semantic properties are not necessarily localized (i.e., “global” attributes, such as color, brightness, orientation or size), which is common in biomedical images. Moreover, interpretability of saliency maps is less informative because they aggregate all of the classification driving image properties to a single heatmap. Counterfactual explanation methods can be subcategorized to those that incorporate latent space disentanglement (such as DISCOVER) and those that do not. Counterfactual explanation methods without disentanglement (Samangouei et al. 2018, Eckstein et al. 2021, Narayanaswamy et al. 2020, Nemirovsky et al. 2020, Shih et al. 2020, Liu et al. 2019, Joshi et al. 2018), can concurrently alter multiple image properties, thus generating less intuitive counterfactual explanations. Counterfactual explanation methods that incorporate disentanglement can be further partitioned to methods that rely on annotated side information of image properties, for example face images with annotated properties such as hair color, mustache, or skin color (He et al. 2019, Gabbay et al. 2021, Li et al. 2020), and to unconditioned methods that do not use any further data annotations beyond the binary classification labels for training the classification model (Lang et al. 2021, Higgins et al. 2021, Rodríguez et al. 2021). DISCOVER benefits from the advantages of both approaches of counterfactual explanations and attribution based methods. Each latent feature is mapped to disentangled classification-driving semantic image properties that can be more intuitively understood by a human observer. DISCOVER does not rely on side annotations, enabling it to discover and quantify unknown subtle semantic image properties which discriminate one class from the other.

Several of DISCOVER’s design choices were proposed by other recent interpretability methods. Several studies included a generator architecture, called “StyleGAN”, that was reported to generate representations that are usually more disentangled than other generative architectures (Wu et al. 2021, Härkönen et al. 2020, Oliva et al. 2020, Lang et al. 2021). Specifically, StylEx uses similar ideas to ours in optimizing latent representations toward high quality counterfactual explanations, along with classification-oriented encoding (Lang et al. 2021). In addition, StylEx instance interpretation relies directly on the latent features values which may suffer from the inherent non-linear associations between latent features and the classifier score. These non-linear associations could hamper latent feature ranking according to their importance toward a specific instance classification prediction. Moreover, all the latent features in StylEx representations are optimized toward all of the model goals, without “specialized” features geared toward specific interpretability goals. In a different study, interpretable directions in the latent space, of a pretrained Generative adversarial network (GAN) generator, were attained by training a new neural network to predict which latent feature was altered to produce a counterfactual explanation in respect to an observed unaltered image (Voynov et al. 2020). DISCOVER’s architecture integrates and extends these ideas. Specifically, our design contributions are (1) disentanglement is explicitly enforced in the latent-to-image space via a new design (loss #5), (2) a focused subset of the latent features is specifically enforced toward classification-driven visual disentanglement (loss #6), (3) direct weighting and ranking of the latent features according to their instance-specific predictive contribution, and interpretation according to the discovered semantic properties that were attributed to each latent feature. Altogether, as we empirically demonstrated in the challenging domain of IVF, these design choices make DISCOVER a designated general-purpose interpretability “discovery machine” especially geared toward quantitative interpretation of known and new classification-driving semantic image properties.

Interpretability of image-based classification models is absolutely necessary in biomedical domains where mechanistic understanding and transparency are crucial. Established attribution-based (Barnett et al. 2021, Kraus et al. 2017, Graziani et al. 2018, Wu et al. 2018, Singh et al. 2020, Zhang et al. 2021) or counterfactual-explanation based (<u>Singla et al. 2023</u>, Thiagarajan et al. 2022, Mertes et al. 2022, Narayanaswamy et al. 2020, Soelistyo et al. 2022, Zaritsky et al. 2021, Lamiable et al. 2023, Kraus et al. 2017) methods were applied, out-of the box or after some adaptations, to interpret a variety of biomedical image-based classification tasks. DISCOVER’s classification-driven and disentanglement representations overcome the inherent limitations in these methods and enabled us to quantitatively confirm non-trivial interpretations, rather than relying on qualitative explanations of representative images, and to systematically perform quantitative instance-specific interpretations.

### Limitations

Although DISCOVER provides a powerful way to uncover the semantic image properties contributing to “black box” classification models’ prediction, it still suffers from several limitations. First, the DISCOVER latent representation is optimized such that each latent feature encodes independent classification-driving semantic image properties. However, this design does not prevent one latent feature to be mapped to multiple independent semantic image properties. In other words, one latent feature may encode entanglement of multiple semantic image properties and still be disentangled in terms of the latent representation. We did not observe examples of 1 (latent feature) - to - many (semantic image properties) in the datasets we explored. Second, DISCOVER may miss semantic image properties that are associated with the classification task. For example, although the inner cell mass (ICM) was a criterion used to define the blastocyst’s quality label, and thus used to optimize the classification model, we failed to interpret a latent feature that encodes the blastocyst’s ICM (Fig. S8). One possible explanation for this inability to interpret the ICM is that other morphological properties may collectively contain the discriminative information encoded in the ICM, and thus, DISCOVER cannot encode the ICM as a classification-driving feature in its latent representation. Indeed, several studies reported that ICM was not an independent predictor of live birth outcome (Ahlström et al. 2011, Hill et al. 2013, Thompson et al. 2013). However, other studies reported that ICM had independent discriminative value (Richter et al 2001, Sivanantham et al. 2022). Another possible explanation is that ICM quality could be explained by combining several more local morphological properties, i.e., it is encoded by multiple classification-driving latent features. Third, we applied DISCOVER to interpret high-performing classification models. Would DISCOVER enable interpretability for less accurate classification models (e.g., Zaritsky et al. 2021)? How well? These are open questions that were not discussed in previous papers, nor here, and will be explored in future studies. We speculate that less accurate classification models will yield more ambiguous visual explanations, thereby making human interpretation less straightforward. Last, we applied DISCOVER to interpret binary classification models. Moving beyond binary classification should be possible by (i) connecting the classification driving subset of latent features to a dense layer of size equal to the number of classes (instead of one neuron) with a softmax (instead of a sigmoid) activation. (ii) changing the classification-driving subset of latent features loss from binary to categorical cross-entropy. (iii) interpreting the classification-driving semantic image properties predictive of a specific class by identifying latent features that correlate with the corresponding softmax probability output. Multi-class interpretability is left for future work.

## Methods

### IVF data collection, annotation and ethics

11,211 embryo time-lapse videos were retrospectively collected from IVF cycles conducted at three clinic centers between March 2010 and December 2021. Historical images of blastocyst-stage embryos and metadata were provided by AIVF LTD. All procedures and protocols were approved by an Institutional Review Board for secondary research use (IRB reference number HMO-006-20). Fertilization (time = 0) was determined by the presence of two pronuclei (2PN) 16-18 hours after insemination. All zygotes were placed inside the EmbryoScope™ time-lapse incubator system (Vitrolife, Denmark), incubated using sequential media protocol until blastocyst-stage, and live imaged with temporal resolution of 15-20 minutes per frame. Each gray-scale image (8bit) was of size 500×500 pixels, with physical pixel size of 294×294 *um*^2^. Z-stacks consisting of 7 slices, 15 µm apart, were acquired at each time point, where the middle slice was used for analysis. Analysis was performed for embryos at the blastocyst stage, with typical onset of blastulation occuring ∼103 hours post insemination based on manual annotation of blastulation and hatching (end of blastulation). 6-10 frames from embryos at the blastocyst stage were collected with an equal time interval between them. High saturated images and images with a partially visible blastocyst were excluded. Overall, approximately 67,000 images were used to train DISCOVER. Blastocysts were manually annotated by embryologists, just before hatching or before the removal of the embryo from the microscope, according to the Gardner and Schoolcraft (known as “Gardner”) scoring criteria, one of the most common morphology-based blastocyst assessment criteria (Gardner et al. 1999). The Gardner criteria is based on three morphology-based quality parameters: (Fig. 1A): Blastocyst expansion status – volume and degree of expansion of the blastocyst cavity (graded 1-6); inner cell mass (ICM) morphology – size and degree of compaction of the mass of cells eventually forming into the fetus (graded A-C); and Trophectoderm (TE) morphology – number and cohesiveness of the single cell layer surround the outer blastocyst eventually forming into the placenta (graded A-C) (Gardner et al. 1998, Gardner et al. 2000). Blastocyst expansion status was not annotated in our dataset. High quality blastocysts were defined by corresponding ICM and TE labels of AA, AB, or BA, low quality blastocysts by BB, BC, or CB.

### Data preprocessing

The image pixel intensities were normalized to the range [0,1]. To accommodate IVF-CLF training on a single GPU (∼30 hours on Nvidia GeForce RTX 3090), the blastocysts images were preprocessed to reduced size, and their background was masked to reduce irrelevant information. Briefly, the preprocessing steps were (1) semantic segmentation of the blastocyst from the raw image, (2) centering the blastocyst in the image, and (3) resizing the image to a lower resolution. Specifically, we trained a mask-RCNN object detection model (He et al. 2017) to detect 200×200 pixels bounding boxes around each blastocyst, using 800 raw images with manually annotated blastocysts’ bounding boxes. Hough-transform (Coste et al. 2012) detected the blastocyst circular shape within the mask-RCNN bounding box and was used to mask the non-blastocyst image regions and to center the blastocyst in the image. Next, a U-NET (Ronneberger et al. 2015) was trained to segment the blastocyst using 500 out of the 800 images that were successfully segmented by the Hough transform (based on manual assessment). The U-NET architecture consisted of 4 convolutional blocks for the encoder (downsampling) with 32, 64, 128 and 256 filters and 4 convolutional blocks for the generator (upsampling) with opposite number of filters. Each convolutional block included a 2D convolution layer, batch normalization and “relu” activation. Max pooling was used for the encoder blocks and upsampling convolution was used for the decoder blocks. The U-NET outputs a binary mask. At inference, the Mask-RCNN is first applied to the raw 500×500 pixels images to output a bounding box localizing the region of the blastocyst. Next, the U-NET uses the localized region and outputs a binary mask, further localizing the blastocyst region. The Hough transform fits a circular contour to the binary mask. This contour mask is multiplied by the Mask-RCNN output to obtain a blastocyst and masked background image. Using the center 2D coordinate of the circular fit, we can center the blastocyst in the image. Finally, the segmented image is resized to 64×64 pixels using nearest neighbors interpolation. The preprocessing pipeline is presented in Fig. S1. Images where the blastocyst was not segmented well (partially cut or large background area remained) were excluded based on visual inspection.

### Classification of high- versus low-quality blastocysts

An ImageNet pretrained VGG-19 network (Simonyan et al. 2014) was fine-tuned by re-training it to discriminate between high-versus low-quality blastocysts (IVF-CLF classifier, Fig. 1C) using a balanced training dataset of 977 high-quality and 977 low-quality blastocysts. Our test dataset was composed of 108 high-quality and 108 low-quality blastocysts. The IVF-CLF architecture is composed of the VGG-19 feature extraction part, which includes several blocks in which each has a downsample convolution layer followed by batch normalization, ReLU activation and a final flatting layer. The last fully connected layer of the pretrained VGG-19 layer (which predicts the 1000 classes of ImageNet) was replaced with a fully connected 16 node dense layer and an output node dense layer with a sigmoid activation, which corresponds to a probability of a high quality blastocyst (0-1). The model was compiled with binary cross entropy loss and Adam optimizer with a learning rate of 0.002. The IVF-CLF network was trained for 100 epochs with a batch size of 32. We performed augmentation by altering brightness, flipping, rotating and by adding Gaussian noise.

### DISentangled COunterfactual Visual interpretER (DISCOVER) architecture and optimization

DISCOVER was designed toward generative interpretability by simultaneously optimizing the following properties (Fig. 2A-B): high-quality and realistic reconstruction of the latent space (loss #1), smooth and realistic traversal of the latent space through its reconstructed images (loss #2), domain-specific classification oriented encoding (loss #3), decorrelated latent space (loss #4), counterfactual disentanglement (loss #5), and a classification-driving subset of latent features that correlated with the classifier that is being interpreted (loss #6). More specifically.

#### Image reconstruction and latent space traversal (losses #1-2)

High-quality and realistic reconstruction and traversal of the latent space was achieved with an adversarial autoencoder (AAE, Makhzani et al. 2015) that was optimized toward a lower dimensional embedded representation of blastocyst images by approximating the high-dimensional data distribution of the input images. This embedding, called latent space, generates a compressed representation that faithfully encodes the input blastocyst. Each blastocyst image is encoded to a point in the latent space that can be decoded to reconstruct an image that appears nearly identical to the original input. The adversarial loss forced the encoded latent representation embedding towards an aggregated posterior distribution similar to a normal distribution in order to achieve a stochastic continuous model to sample from during traversal (Makhzani et al. 2015). The encoder (Table S1) and decoder (Table S2) networks backbone were based on residual blocks similar to the ones introduced in Resnet50 (He et al. 2016). The outputs of the last convolutional downsampling block were flattened to a vector followed by a dense layer of 350 dimensions (determined empirically) that defined the latent representation. The discriminator network was composed of six fully connected dense layers (Table S3).

The reconstruction loss (loss #1) was a perceptual loss where the reconstruction minimized the Euclidean distance between the hidden layers of a VGG-19 pre-trained on ImageNet (called Imagenet-CLF). Perceptual loss was preferred over minimizing L1 or L2 pixel-wise differences because the latter lead to blurry, and less realistic reconstructed images (Fig. S4A). Perceptual loss enforces spatial consistency between the real and the reconstructed images which is important for human interpretability (Pihlgren et al. 2020, Zhang et al. 2018). More technically, for a blastocyst image x, and its corresponding reconstructed image x_rec_, we extracted the hidden representations of the Imagenet-CLF network: Imagenet-CLF(x)^i^ and Imagenet-CLF(x_rec_)^i^ from layers i =[block3_conv1, block3_conv2, block3_conv3, block4_conv1, block4_conv2, block4_conv3, block4_conv4, block5_conv1, block5_conv2, block5_conv3, block5_conv4]. For every layer the mean absolute error (MAE) was calculated and the overall losses was an average of these per-layer (i) errors:

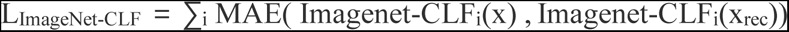

The latent generative-adversarial loss (loss #2) enforced a probabilistic latent space such that samples were encoded into a continuous dense distribution. Adversarial losses are designed to fool a discriminator: a discriminator network (D) is trained to predict if an input vector comes from the latent representation of the encoded images z, or drawn from the normal distribution with mean 0 and variance of 1, z_noise_. The adversarial loss pushes the encoder to output latent representations with a similar normal distribution. The discriminator receives either the encoderoutput z or a noise vector z_noise_ and predicts the source (encoded versus noise). The discriminator loss is a binary cross-entropy loss:

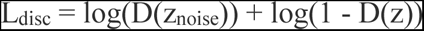

and the encoder (E) adversarial loss is:

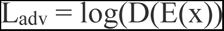

#### Classification oriented encoding (loss #3)

Subtle image differences can lead to major differences in the classification outcome. Thus, to ensure that the visual semantic properties influencing the classification decision are maintained in the reconstructed image, we introduced a loss term that minimized the discrepancy between the IVF-CLF hidden representations of the real versus its corresponding reconstructed image (Fig. 2A). Similarly to loss #1, we minimized the perceptual loss by extracting the hidden representations of the IVF-CLF network: IVF-CLF(x)^i^ and IVF-CLF(x_rec_)^i^ from layers i = [block3_conv1, block3_conv2, block3_conv3, block4_conv1, block4_conv2, block4_conv3, block4_conv4, block5_conv1, block5_conv2, block5_conv3, block5_conv4, flatten, dense]. For every layer the mean absolute error (MAE) was calculated and the overall losses was an average of these per-layer (i) errors:

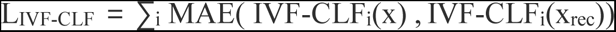

#### Disentangled latent representation (losses #4-5)

The disentanglement module (Fig. 2B) was designed to encode the image into a decorrelated latent space, where each latent feature is independent (i.e., decorrelated) from the others (loss #4) and is associated with a distinct visual property in the image (loss #5).

A decorrelated latent representation encourages each latent feature to be independent of other latent features. We included a loss which whitens the latent features’ covariance matrix (i.e., driving it to become a unit matrix), by optimizing toward diagonal values of 1 and off-diagonal values to 0, similar to (Bardes et al. 2021):

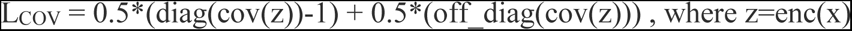

Disentanglement of the latent representation enables traversal of the latent space one feature at a time under the assumption that each latent feature encodes an independent classification-driving visual image features. To enforce that a specific latent feature is associated with a specific image property we minimized the error of an additional neural network that was trained to identify which latent feature was altered upon alteration of a single latent feature. This was implemented by (1) altering a randomly selected latent feature value in the range of ±1.5 standard deviations, (2) using the decoder to reconstruct a blastocyst image from the altered latent vector, (3) constructing a “diff image”, the subtraction of the altered reconstructed image from the unaltered reconstructed image, (4) A disentanglement network (Table S4) is trained to predict the index of the latent feature that was altered from an input of the “diff image”. The disentanglement network was implemented by down sampling convolutions followed by a flattening layer and a dense layer equal to the size of the latent space (Nz = 350) along with a Softmax activation, and outputs a latent feature probability. Categorical cross-entropy (CCE) loss was used to minimize the difference between the output prediction vectors of the network after softmax activation ypred and the one-hot encoding vector ytrue, where the altered latent feature value was set to 1 and all other latent features were set to a value of 0:

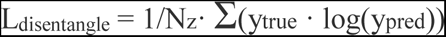

Note that the backpropagation of this loss term goes all the way back through the decoder and encoder, thus enforcing visual disentanglement as an inherent property of the latent representation.

#### Classification-driving subset of latent features (loss #6)

We designed a loss to partition the latent representation to two subsets: (1) 14 latent features that are correlated to the IVF-CLF classification score, i.e., associated with semantic properties driving the classifier’s decision; (2) The other 336 latent features maintain high-quality reconstruction without enforcing correlation to the IVF-CLF score. We call the first subset “classification-driving”, and the latent features in this subset can be altered to create reconstructed blastocysts images with corresponding alteration in the IVF-CLF classification output and thus can be used toward interpretation of the semantic classification-driving physical properties that they encode. The size of the classification driving subset was determined under the assumption that a small subset would be more interpretable. The counterfactual disentanglement network was implemented as a single neuron trained to predict the IVF-CLF’s classification score from the classification driving subset, which were the first 14 features in the latent representation. The latent features in classification driving subset were connected via a dense layer to the single classification neuron with a sigmoid activation (Z_subset_score_), and the binary cross entropy (BCE) between the prediction and the IVF-CLF scores was minimized.

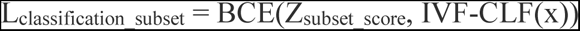

#### Optimization

The necessity of all loss terms was verified via ablation experiments (Fig. S4). The overall loss of DISCOVER was defined as the addition of all six loss terms, with weights λ_1_=5, λ_2_=1, λ_3_=5, λ_4_=1, λ_5_=1, λ_6_=1, that were adjusted empirically by observing that the reconstruction losses were converging slower than other losses. Thus, the following loss was minimized during training:

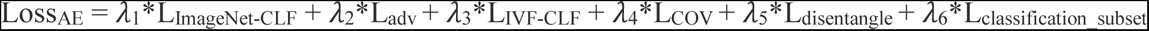

DISCOVER was trained with Adam optimizer, learning rate of 0.0002 and batch size of 64. It was trained for 30 epochs. In each iteration, images were chosen randomly and the following augmentations were performed for the IVF dataset: (1) brightness - randomly multiplying each image by a factor of −0.2 to 0.2. (2) flip - randomly flipping images horizontally and vertically. (3) rotation - randomly rotating images by 0, 90, 180, or 270 degrees. (4) noise - introducing per-pixel Gaussian noise was added with mean 0 and standard deviation of 0.1. (5) saturation - random pixels’ gray levels were saturated.

### Visualization of counterfactual alteration

Counterfactual alterations, the changes in image properties associated with the change of a latent feature, were visualized using the Structural Similarity Index (SSIM) (Renieblas et al. 2018). SSIM has been demonstrated to be in agreement with how humans observe differences between two images. SSIM evaluates the similarity of two images by comparing spatially matched pairs of image patches using the average, standard deviation and covariance of each patch. For visualization, each pixel was assigned the value 1-SSIM, corresponding to the dissimilarity between the two corresponding patches of 7×7 pixels surrounding the pixel. This was followed by smoothing with a convolution with a gaussian filter of size 3×3 to define what we call the “*visual counterfactual alteration*“.

### Quantitatively validating latent features interpretation

To systematically and quantitatively link latent features #0 and #10 to their corresponding interpreted morphological properties, we had to reduce the confounding effect of the correlated blastocysts’ size and TE. Thus, we matched pairs of blastocysts according to having one similar morphological property, and the other morphological property being different. More specifically, to verify that latent feature #0 is associated with blastocyst size, we paired blastocysts according to (i) same embryologist-annotated TE grades, i.e., both blastocysts with grade ‘A’ or both with grade ‘B’; (ii) at least 30% difference in their sizes, i.e., the size of the larger blastocyst was ≥ 1.3 times of the smaller blastocyst. Blastocyst size was computed from the segmented blastocyst masks as described earlier (see the subsection “Data preprocessing”). A total of 5,888 blastocysts’ pairs were matched according to these criteria. To measure the association between each of the 14 classification-driving subsets of latent features and the blastocyst size, we calculated the distribution of signed differences of each latent feature between the larger and the smaller blastocysts in the matched pairs (Fig. 4A). Importantly, the subtraction order was flipped for latent features that were negatively correlated with the IVF-CLF scores (Fig. 2F). The purpose of adjusting the subtraction order according to the correlation sign was to enable direct ranking ordering of the associations between the latent features and the blastocyst size, for matched blastocysts (with the same TE annotations), according to the median of each (latent feature specific) signed differences distribution (Fig. 4B-C). The direct comparison between distributions was enabled by z-score normalization of the latent features.

Similar analysis was performed to verify that latent feature #10 was associated with the blastocyst TE quality. Blastocysts were paired according to (i) different embryologist-annotated TE grades, i.e., one blastocysts with grade ‘A’ and the other with grade ‘B’; (ii) no more than 7% difference in their sizes. A total of 808,326 blastocysts’ pairs were matched according to these criteria. Similarly to the analysis that linked latent feature #0 to blastocyst size, we measured the association of each of the 14 classification-driving subsets of latent features and the blastocyst TE quality, where the order of subtraction was determined according to the sign of the correlation between the latent feature values and the IVF-CLF scores.

### Instance interpretation

To quantify the latent features importance to the classification prediction we used Shapley additive explanation (SHAP) (Lundberg et al. 2017). We applied SHAP on DISCOVER’s single layer perceptron which receives as input the 14 classification-driving subset of latent features and is connected to a single neuron upon which a sigmoid activation is applied to predict the IVF-CLF score (Fig. 6A). The estimated average SHAP values for each latent feature was calculated using a random subset of 200 samples (Fig. S6).

### Embryologists qualitative feedback and quantitative validations

Embryologists provided qualitative feedback and participated in a user-study to quantitatively validate our interpretations. For qualitative feedback of GradCAM’s interpretability, two embryologists were presented with visual explanations of 18 blastocysts (those shown Fig. 1E) obtained by GradCAM, highlighting the important localized regions of the IVF-CLF’s final convolutional block. The embryologists were asked whether the GradCAM visualizations provide insight regarding the blastocyst’s morphological properties that were learned by the model. For qualitative feedback of DISCOVER’s disentanglement and interpretability, two embryologists were presented with counterfactual visual alterations of the same blastocyst according to the alteration (± 3 standard deviations) of the five latent features most correlated to the IVF-CLF (see example in Fig. 3B, this evaluation was performed for 3 blastocysts). The embryologists were asked to interpret the morphology that changed between the counterfactual explanations for each of the latent features. To qualitatively validate our interpretation of latent features #0 and #10 as encoding the blastocyst size and TE, respectively, two Embryologists were (i) presented with the counterfactual visual alterations of 16 blastocysts (Fig. S5), (ii) presented with a sequence of gradually altered traversals (± 3 standard deviations) along each latent feature (Fig. 3C), (iii) presented with a sequence of nine blastocysts randomly selected and ordered according to their corresponding latent feature values, in equal intervals along the range of ± 3 standard deviations, for latent features #0 and #10 (Fig. 3D). For each of these evaluations, the embryologists were asked to describe which visual property was mostly dominant. Latent feature #11 was interpreted and qualitatively validated to be associated with the blastocoel density by presenting to a trained embryologist and two other IVF experts (i) counterfactual visual alterations of 5 blastocysts (Fig. 5B), (ii) a sequence of 9 real blastocysts that were randomly selected from predefined intervals of i latent feature #11 in monotonically increasing order (Fig. 5C). To quantitatively verify that latent features #0 and #10 encode the blastocyst size and TE, respectively, we performed an empirical user study. For the user study we matched 39 blastocyst pairs according to a similar value (< 0.1) of latent feature #0, and a different value (> 0.5) of latent feature #10. values, and 36 blastocyst pairs according to a similar value (< 0.05) of latent feature #10 and different value (> 0.6) of latent feature #0 values. The different thresholds for “similar” or “different” were selected to achieve a close number of blastocyst pairs selected according to each of the two conditions. These 75 blastocyst pairs were presented, in a random order, to an embryologist that was asked to determine which morphology (size or TE) was more different between the two blastocysts in each pair. Additionally, the embryologist was asked to determine which blastocyst within each pair had a higher grade of that dominant morphology. A confusion matrix and accuracy results of our user study are reported in Fig. S3.

To qualitatively verify the interpretation of specific blastocysts’ classification (see the subsection “Instance interpretation”), three high quality and three low quality blastocysts were randomly selected according to the following criteria: two with SHAP-dominating latent feature #0 (Fig. 6B left), two with SHAP-dominating latent feature #10 (Fig. 6B middle), and two with SHAP-dominating latent features #0 and #10 (Fig. 6B right). These six blastocysts were presented to an embryologist who visually verified the instance-specific SHAP feature importance according to our mapped interpretation (latent feature #0/#10 encode size/TE). Similarly, three blastocysts composed of two positive and one negative SHAPE-dominating latent feature #11 were randomly selected and visually verified the instance-specific SHAP feature importance according to our mapped interpretation (blastocoel) by the embryologist.

### CelebA faces dataset, preprocessing, gender classification, and DISCOVER interpretability

The celebA dataset (Liu et al. 2015) contains 202,599 aligned and cropped RGB images (64×64 pixels) of 10,000 celebrities’ faces with an associated male/female attribute (as well as additional 40 binary annotations such as smile, hat etc.). We trimmed 15 pixels from each side to remove background nuisance and the image was then resized back to 64×64 pixels. All images were converted to grayscale and divided by 255 to the range [0-1]. A VGG-19 classification model was trained to discriminate between male and female face images, we call this model GENDER-CLF. The training followed the same procedure described for IVF-CLF (see Fig. 1C, and earlier in the Methods). 15,000 images from each gender were randomly selected for training. The AUC for the test data (1,000 images for each gender) was 0.96 (Fig. S7A). No augmentations were used in this training. We trained a DISCOVER network to interpret GENDER-CLF. The differences in training, in respect to interpreting IVF-CLF were a training set composed of 164,268 (65,183 male and 99,085 female) images without augmentations.

### Statistical analysis

ROC-AUC (sklearn.metrics.auc function) was used to evaluate the performance of the classifier models (Fig. 1D, Fig S7). Pearson correlation (scipy.stats.pearsonr function) was used to assess the inner correlations between the latent features (Fig. S2) and the correlation between each latent feature and classifier score (Fig. 2F). Mann–Whitney-U test (scipy.stats.mannwhitneyu) was used to calculate the p-value of the 14 classification-driving subset of latent features out of the entire latent feature representation.

### Code and data availability

The source code (Python with Tensorflow 2.2) for training a binary classifier, training a DISCOVER interpretability model and a demonstration of performing blastocyst classification interpretability using a trained DISCOVER model are publicly available, https://github.com/OdedRotem314/DISCOVER. We are currently working toward contributing these models to the Bioimage Model Zoo to make them more accessible (<u>Ouyang, Beuttenmueller, Gómez-de-Mariscal, et al. 2022</u>).

This repository also includes a trained model for gender classification and its corresponding DISCOVER model. The celebA dataset is available https://mmlab.ie.cuhk.edu.hk/projects/CelebA.html.

## Funding and Acknowledgements

This research was supported by the Israel Council for Higher Education (CHE) via the Data Science Research Center, Ben-Gurion University of the Negev, Israel (to AZ), and by the Rosetrees Trust (to AZ). We thank Lion Ben Nedava, Alon Shpigler, Lior Rokach, Meghan Driscoll, and Orit Kliper-Gross for critically reading the manuscript. We dedicate this manuscript to Jonathan Seidman, son of our co-author, Daniel S. Seidman, who was brutally murdered during a tragic terrorist attack perpetrated by Palestinian terrorists, while participating in a Peace Music Festival. This tragedy has left DSS, the rest of the authors, and our nation profoundly devastated. This manuscript was finalized in the wake of these events, while we grieve and mourn.

## Author Contribution

OR and AZ conceived the study. OR developed the computational tools, and analyzed the data. TS annotated data. TS, MTS and RN confirmed visual interpretation. OR and AZ interpreted the data and drafted the manuscript. RM, YT, MTS, MM, DG and DSS provided clinical input for the presentation of the manuscript, reviewed and revised the manuscript. AZ mentored OR. All authors edited the manuscript and approved its content.

