## Supplemental figures and tables for "Visual interpretability of image-based classification models by generative latent space disentanglement applied to in vitro fertilization"

### Supplemental figures

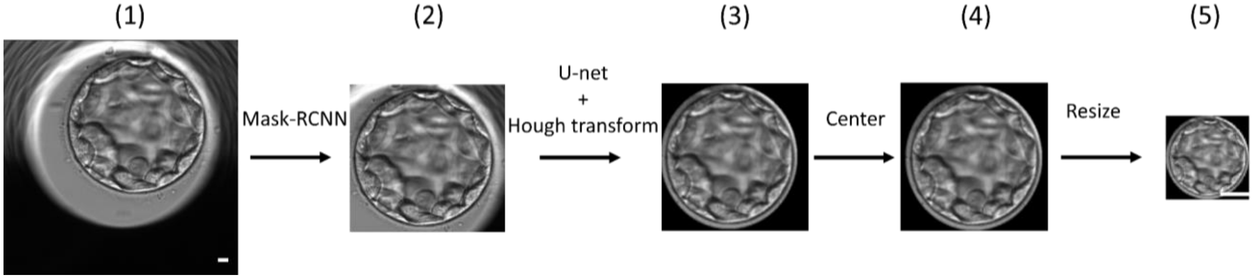


**Figure S1. Preprocessing (left-to-right)**. (1) A snapshot blastocyst image (500x500 pixels = 294.1x294.1$\mu m^2$) from the time-lapse sequence at 103 hours past insemination. (2) The blastocyst region was localized with a bounding box of 200x200 pixels obtained by a Mask-RCNN object detection model. (3) The blastocyst was segmented from the background by a U-NET segmentation model followed by Hough transform and the background pixels were assigned to 0 (black). (4) The blastocyst was centered in the bounding box according to the circular fit obtained by the Hough transform. (5) The bounding box image was rescaled to 64x64 pixels. Scale bar = 12.5 $\mu$m.

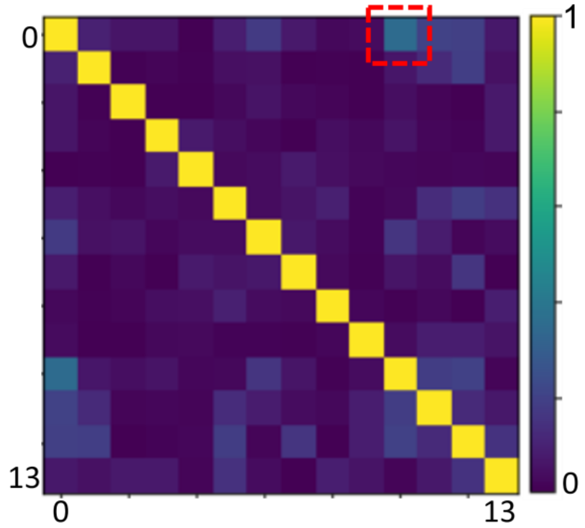


**Figure S2. Pairwise latent feature correlation**. Pairwise absolute Pearson correlation coefficient between each pair of the 14 classification-driving subset of latent features. The correlation between latent features #0 and #10 is weak (-0.35) but also the most prominent and is highlighted by the red dashed square.

| | | |
|---|---|---|
| True size 30 | False size 6 | PPV 0.83 |
| False TE 4 | True TE 35 | NPV 0.1 |
| Sensitivity 0.88 | Specificity 0.14 | Accuracy 0.86 |

**Figure S3. Confusion matrix of embryologist validation**. 75 matched random blastocyst pairs were selected with either different values (> 0.6) of latent feature #0 and similar (< 0.1) values of latent feature #10 or vice-versa (i.e., similar #0, different #10). The embryologist was asked to determine which property (size/TE) was different from the corresponding blastocysts images. “True size” - different feature #0 and the embryologist answered “size”, “True TE” - different feature #10 and the embryologist answered “TE”, “False size” - different feature #0 and the embryologist answered “TE”, “False TE” - different feature #10 and the embryologist answered “size”. For “True size” pairs the embryologist’s answer to which blastocyst had larger size (higher feature #0 value) was 31/36 (86%). For “True TE” pairs the embryologist’s answer to which blastocyst had better TE quality (lower feature #10 value) was 33/39 (85%).

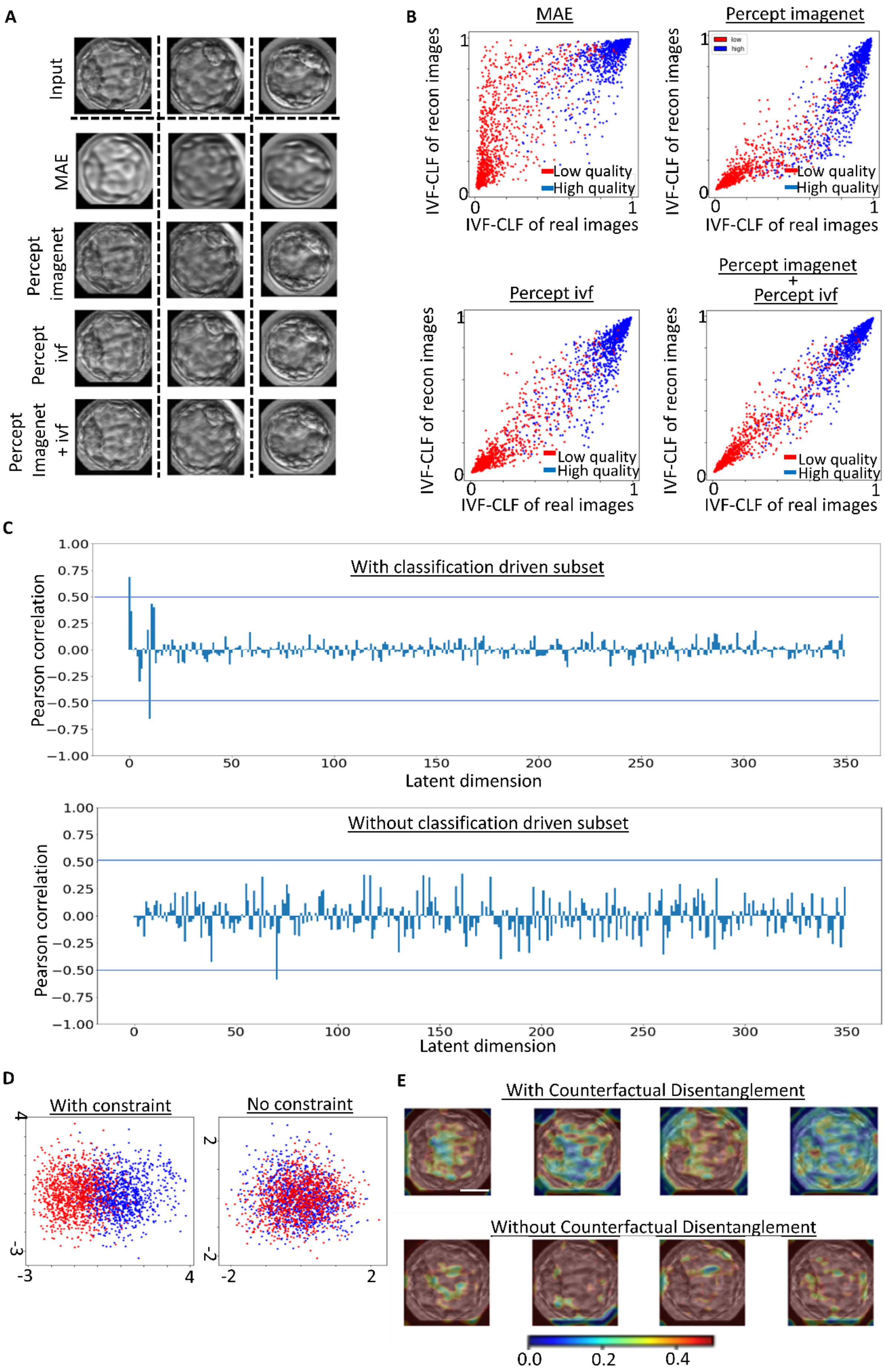
A
Input
MAE
Percept imagenet
Percept ivf
Percept Imagenet + ivf
B
MAE
Percept imagenet
Percept ivf
Percept imagenet + Percept ivf
IVF-CLF of recon images
IVF-CLF of real images
Low quality
High quality
low
high
C
With classification driven subset
Without classification driven subset
Pearson correlation
Latent dimension
D
With constraint
No constraint
E
With Counterfactual Disentanglement
Without Counterfactual Disentanglement
0.0
0.2
0.4

**Figure S4. Ablation study**. (**A-B**) High-quality image reconstruction (loss #1) using a classification oriented encoding (loss #3). Qualitative visual similarity (A) and quantitative similarity in IVF-CLF classification scores (B) compared across four reconstruction losses: (i) mean absolute error (MAE) - pixel-wise L1 distance; (ii) ImageNet-CLF perceptual loss (percept imagenet); (iii) IVF-CLF perceptual loss (percept ivf) (iv) ImageNet-CLF perceptual loss and IVF-CLF perceptual loss (percept imagenet + percept ivf). (**A**) Reconstruction of three representative blastocyst images (columns) according to the four losses (rows). Scale bar = 12.5 $\mu$m. (**B**) Scatter plots showing the correlations between the IVF-CLF classification score of the blastocysts' images (x-axis) and their matched reconstructed images according to the four losses (y-axis). N = 2169 blastocysts that were not used to train the model, marked in blue (high quality) or red (low quality) blastocysts. The mean absolute error between real images scores and reconstructed images scores was 0.14 (MAE), 0.10 (percept imagenet), 0.07 (percept ivf), and 0.04 (percept imagenet + percept ivf). (**C-D**) Classification-driving subset of latent features (loss #6). (**C**) Pearson correlation coefficient (y-axis) between all 350 latent features (x-axis) and the IVF-CLF's classification score with (top, duplicate of Fig. 2F) or without (bottom) loss #6. Same blastocysts as in B. **(D)** Principal components analysis (PCA) of the 14 classification-driving subsets of latent features, i.e., with loss #6 (left) versus the 14 most correlated features without loss #6 (right). Shown PC1 (x-axis) and PC2 (y-axis) the two top principal components encoding 0.099 and 0.092 (without loss #6) and 0.13 and 0.09 (with loss #6). Inclusion of loss #6 separates better the high quality (blue) and the low quality (red) blastocysts. Same blastocysts as in B. (**E**) Counterfactual disentanglement of the latent representation (loss #5). SSIM visualization (see text and Methods) of counterfactual alteration of the same blastocyst. Each column from left to right represents the ranked latent feature correlation to the IVF-CLF (left being the strongest correlation and with loss #5 (top row) or without (bottom row) loss #5. Note distinct (top) versus aggregate (bottom) visual counterfactual alteration patterns with and without the counterfactual disentanglement loss term correspondingly. Scale bar = 12.5 $\mu$m.

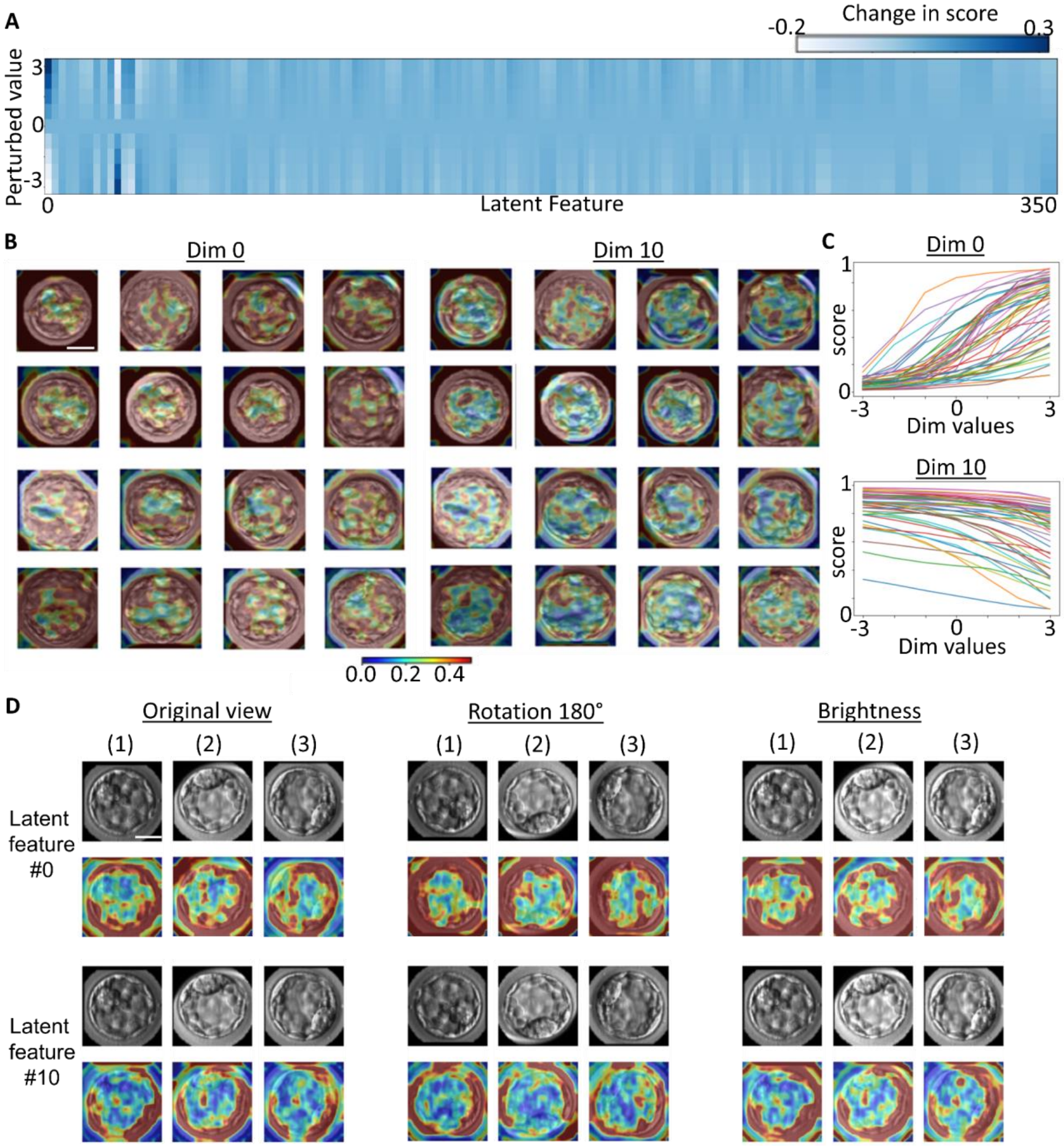


**Figure S5. Classification-associated latent feature #0 encodes the blastocysts' size, latent feature #10 encodes the trophectoderm quality.** (**A**) Change in IVF-CLF scores upon perturbing each latent feature 0 ± 3 standard deviations. Reported the mean change in classification score of the reconstructed blastocyst image before and after alteration of each latent feature, averaged across N = 2169 blastocysts. The classification-driving subset of latent features, and especially features #0 and #10, exhibit higher change in the classification score in response to their alteration. (**B**) Representative blastocysts' counterfactual visual alterations, by ± 3 standard deviations, of latent feature #0 encodes the blastocyst size (left), and latent feature #10 encodes the trophectoderm (right). **(C)** Gradual monotonic change in the IVF-CLF classifier score of 30 random blastocysts reconstructions after gradual traversal along latent feature #0 (left) and #10 (right). **(D)** Robust visual interpretability to rotation of 180 degrees (middle panel) and multiplying the image by a brightness factor of 1.15 (bottom panel) in respect to the raw images (top panel). Each panel shows the reconstruction of the same 3 representative blastocysts (columns 1-3) ± 3 standard deviations in the corresponding latent feature (#0, #10) from the observed blastocyst image encoding. Scale bar = 12.5 $\mu$m.

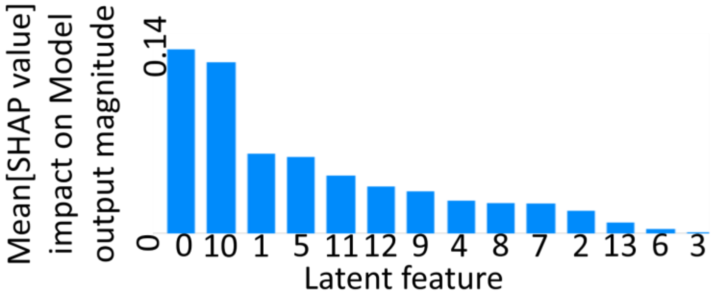


**Figure S6. Mean latent features SHAP**. Mean SHAP values (y-axis) of the 14 classification-driving subsets of latent features (x-axis).

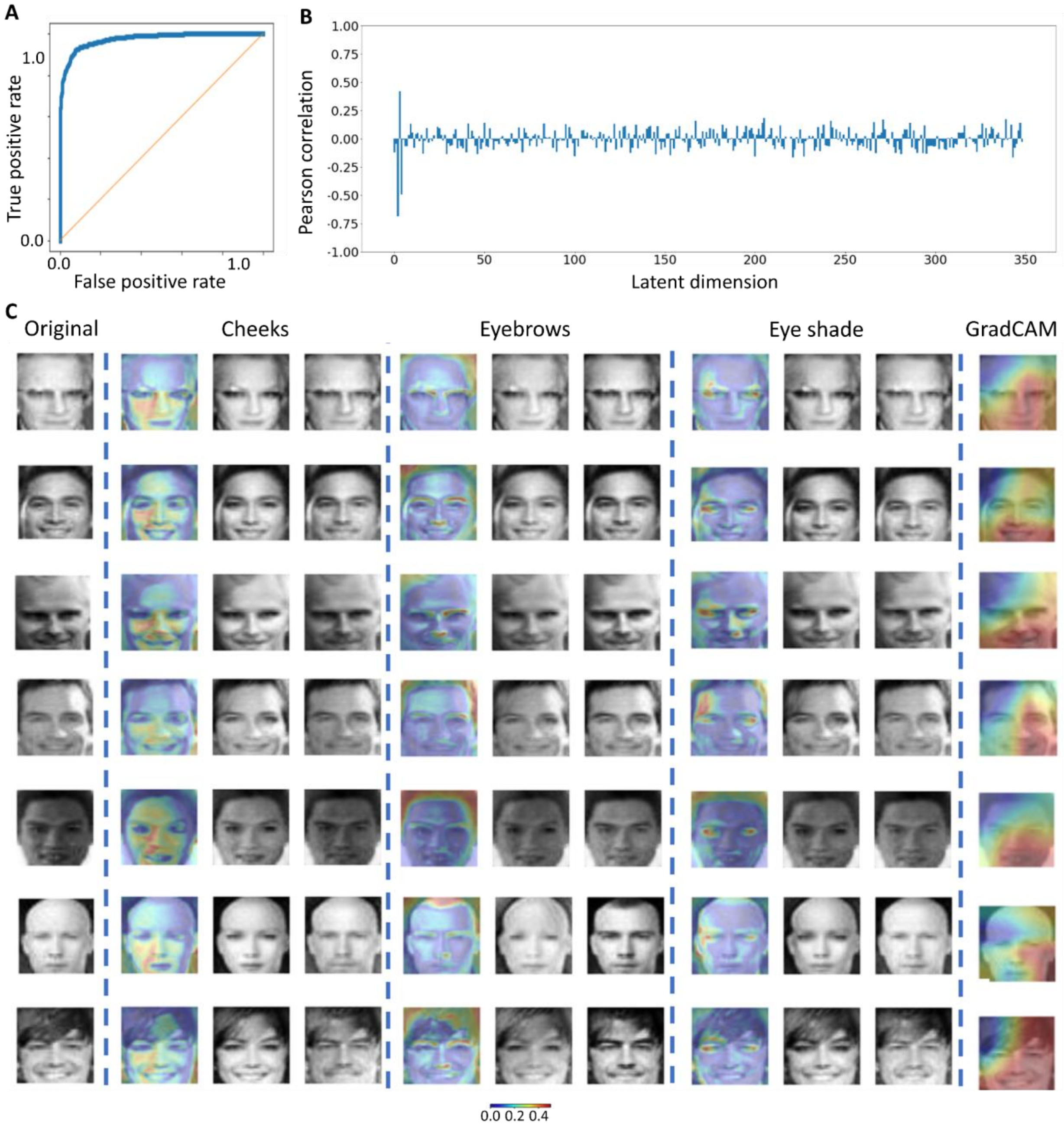


**Figure S7. Learning the facial features which reveal the differences between male and female**. Data from the public CelebA dataset, https://mmlab.ie.cuhk.edu.hk/projects/CelebA.html, posting these photos do not raise any conflict with the copyright policy of the database. (**A**) ROC curve of the GENDER-CLF with train/test sets of 15K/1K for every gender, test AUC 0.96. (**B**) Pearson correlation coefficient (y-axis) between each latent feature (x-axis) and the GENDER-CLF's classification score, using a test set of 2K faces that were not used to train DISCOVER (trained with 164,268 images). (**C**) seven examples showing the three main facial features which alter the gender of the face. First column presents the original image. Second, third and fourth main columns (separated by dashed lines) correspond to latent features #2, #4 and #3 with Pearson correlation coefficient of 0.68, 0.49, and 0.42, respectively. correspondingly Each column shows the visual counterfactual alteration map (left colored image), female altered image and male altered image when traversing the latent feature by +3 or -3 std (if the correlation sign is negative then vice versa). The last column shows the Gradcam heatmap for each example.

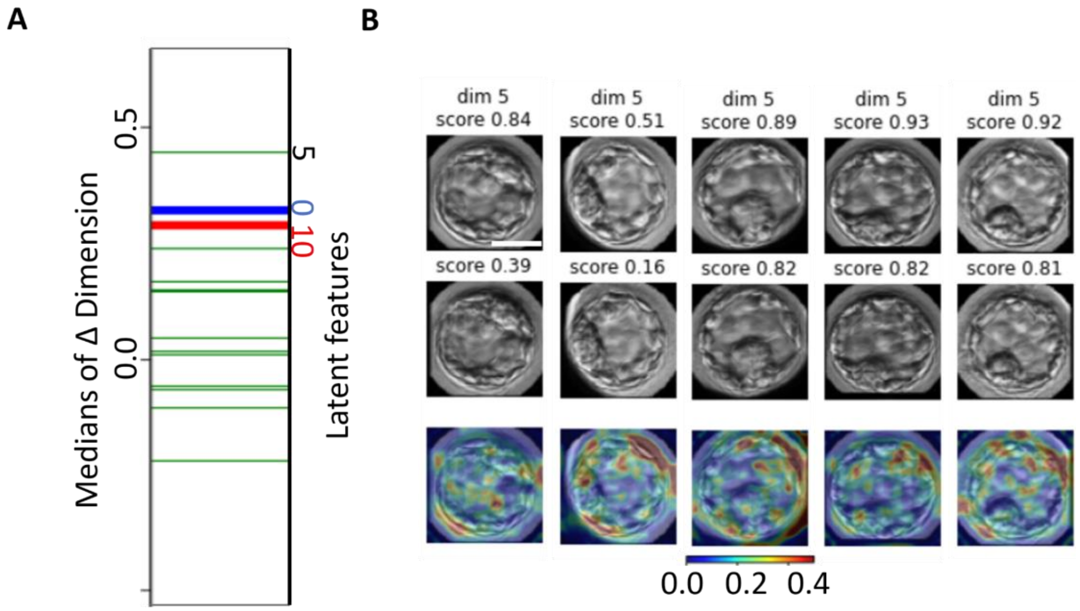


**Figure S8. Unsuccessful attempt to interpret a latent feature encoding the blastocyst's inner cell mass (ICM)**. **(A)** Analysis similar to Fig. 4. 2,134 pairs of blastocysts matched according to having similar TE (both 'A' or both 'B'), similar size (size difference < 0.007%) and different ICM ('A' with 'B' or vice versa). The subtractions of a latent feature value of one blastocyst and its corresponding paired blastocysts were pooled for each latent feature. The order to subtraction was determined according to the sign of the correlation between the latent feature and the IVF-CLF scores (Fig. 2F). The median values of the distributions of signed differences for each of the 14 classification-driving subsets of latent features are shown. The blue and red vertical line represent the median of latent features #0 and #10 respectively. Latent feature #5 was ranked first suggesting that it specifically encodes the ICM morphology. (**B**) Counterfactual visual alteration of five blastocysts obtained by altering latent feature #5 by ± 3 standard deviations. Top row: reconstructed altered images with increased classification score. Middle row: reconstructed altered images with reduced classification scores. Bottom row: counterfactual visual alteration between the corresponding top and middle rows. Color map indicates the local change measured as 1-SSIM. Scale bar = 12.5 $\mu$m.

### Supplemental tables

Each table describes the architecture details of the network. Details include each layer type and its corresponding input feature maps dimensions, output feature maps dimensions, filter size, stride, using spectral norm, using batch norm and which activation function.

Resblock structure:

Input⇒BN⇒Relu⇒Conv2D(filters, kernel=3, stride=2) ⇒ BN⇒Relu⇒Conv2D(filters, kernel=3, stride=1) which is added to Input⇒Conv2D(filters, kernel=1, stride=2)

| Layer | Input | Output | Filter size | Stride | Spectral norm | Batch Norm | Activation |
|---|---|---|---|---|---|---|---|
| Conv2D | (64,64,1) | (64,64,64) | 3x3 | 1 | Yes | No | ReLU |
| Resblock Down sample | (64,64,64) | (512,32,32) | 3x3 | 2 | Yes | Yes | ReLU |
| Resblock Down sample | (512,32,32) | (512,16,16) | 3x3 | 2 | Yes | Yes | ReLU |
| Resblock Down sample | (1024,16,16) | (1024,8,8) | 3x3 | 2 | Yes | Yes | ReLU |
| Resblock Down sample | (1024,8,8) | (1024,4,4) | 3x3 | 2 | Yes | Yes | ReLU |
| Flatten | (1024,4,4) | (16,384,) | | | No | No | Swish |
| Dense | (16,384,) | (350,) | | | No | No | Swish |
| Lambda split | (350, ) | (10,) | | | | | |
| Dense | (10,) | (1,) | | | No | No | Sigmoid |

Table S1: Encoder architecture

| Layer | Input | Output | Filter size | Stride | Spectral norm | Batch Norm | Activation |
|---|---|---|---|---|---|---|---|
| Dense | (350,) | (16,384,) | | | No | No | |
| Reshape | (16,384,) | (1024,4,4) | | | | | |
| Resblock Up sample | (1024,4,4) | (1024,8,8) | 3x3 | 2 | Yes | Yes | ReLU |
| Resblock Up sample | (1024,8,8) | (1024,16,16) | 3x3 | 2 | Yes | Yes | ReLU |
| Resblock Up sample | (1024,16,16) | (512,32,32) | 3x3 | 2 | Yes | Yes | ReLU |
| Resblock Up sample | (512,32,32) | (512,64,64) | 3x3 | 2 | Yes | Yes | ReLU |
| Conv2D | (512,64,64) | (1,64,64) | 3x3 | 2 | Yes | Yes | Sigmoid |

Table S2: Decoder architecture

| Layer | Input | Output | Filter size | Stride | Spectral norm | Batch Norm | Activation |
|---|---|---|---|---|---|---|---|
| Dense | (350,) | -2048 | | | No | Yes | LeakyReLU |
| Dense | (2048,) | -2048 | | | No | Yes | LeakyReLU |
| Dense | (2048,) | -2048 | | | No | Yes | LeakyReLU |

| | | | | | | | |
|---|---|---|---|---|---|---|---|
| Dense | (2048,) | -2048 | | | No | Yes | LeakyReLU |
| Dense | (2048,) | -2048 | | | No | Yes | LeakyReLU |
| Dense | (2048,) | -2048 | | | No | Yes | LeakyReLU |
| Dense | (2048,) | -1 | | | No | Yes | Sigmoid |

Table S3: Discriminator architecture

| **Layer** | **Input** | **Output** | **Filter size** | **Stride** | **Spectral norm** | **Batch Norm** | **Activation** |
|---|---|---|---|---|---|---|---|
| Conv2D | (64,64,1) | (64,64,64) | 3x3 | 1 | Yes | No | ReLU |
| Resblock Down sample | (64,64,64) | (512,32,32) | 3x3 | 2 | Yes | Yes | ReLU |
| Resblock Down sample | (512,32,32) | (512,16,16) | 3x3 | 2 | Yes | Yes | ReLU |
| Resblock Down sample | (1024,16,16) | (1024,8,8) | 3x3 | 2 | Yes | Yes | ReLU |
| Resblock Down sample | (1024,8,8) | (1024,4,4) | 3x3 | 2 | Yes | Yes | ReLU |
| Flatten | (1024,4,4) | (16,384,) | | | No | No | Swish |
| Dense | (16,384,) | (350,) | | | No | No | Softmax |

Table S4: Disentangler architecture
